# Cardiac hemodynamics computational modeling including chordae tendineae, papillaries, and valves dynamics

**DOI:** 10.1101/2024.05.21.595150

**Authors:** Andrea Crispino, Lorenzo Bennati, Christian Vergara

## Abstract

In the context of dynamic image-based computational fluid dynamics (DIB-CFD) modeling of cardiac system, the role of sub-valvular apparatus (chordae tendineae and papillary muscles) and the effects of different mitral valve (MV) opening/closure dynamics, have not been systemically determined. To provide a partial filling of this gap, in this study we performed DIB-CFD numerical experiments in the left ventricle, left atrium and aortic root, with the aim of highlighting the influence on the numerical results of two specific modeling scenarios: i) the presence of the sub-valvular apparatus, consisting of chordae tendineae and papillary muscles; ii) different MV dynamics models accounting for different use of leaflet reconstruction from imaging.

This is performed for one healthy and one MV regurgitant subjects. Specifically, a systolic wall motion is reconstructed from time-resolved Cine-MRI images and imposed as boundary condition for the CFD numerical simulation.

Analyzing the numerical results, we found that sub-valvular apparatus do not affect the global fluid dynamics quantities, although it creates local variations, such as the developing of vortexes or flow disturbances, which lead to different stress distributions on cardiac structures.

Moreover, different MV dynamics are considered starting from Cine-MRI MV segmentation at different temporal configurations, and then they are compared and managed numerically through a resistive approach. The obtained results highlight the importance of including a sophisticated diastolic model of MV dynamics, which accounts for MV geometries during diastasis and A-wave, in terms of describing the disturbed flow and ventricular turbulence.

**Statements and Declarations:** The authors have no relevant financial or non-financial interests to disclose.

## 1 Introduction

The study of the ventricular hemodynamics plays a key role in understanding physical and physiological cardiac phenomena, from a global to a local level. A very useful tool for carrying out this study consists in the use of cardiac computational models, that can be distinguished into three main categories: *electro-fluid-structure interaction* (EFSI) models [94, 14], *fluid-structure interaction* (FSI) models with prescribed or surrogate electro-physiology [59, 33, 83, 34, 88, 26, 16, 27, 17, 89, 28], and *computational fluid dynamics (CFD) with prescribed motion* of the heart walls. Regarding this latter approach, the required cardiac motion can be obtained through either an *electromechanic* simulation [98, 6, 45, 102] or by processing and employing dynamic (time-resolved) medical images (*Dynamic Image-based CFD*, DIB-CFD) [8, 51, 19, 84, 62, 32]. In this context, a crucial role in view of the assessment of accurate results is played by the modeling of the *sub-valvular mitral apparatus* and of the *opening/closure dynamics* of the mitral valve (MV).

The sub-valvular apparatus consists on *papillary muscles* and *chordae tendineae* (Figure 1). Papillary muscles (PM) are small myocardial structures, located within the cavity of the left ventricle (LV), attached to its walls; according to their position, they can be divided into anterolateral PM and postero-medial PM [60]. Chordae tendineae (CT) are fan-shaped strings of fibrous connective tissue that run from the tip of PM and inserts into different regions of mitral valve leaflets [79]; depending on the region of attachment, different types of chordae can be defined, according to Lam?s functional classification [50]. PM and CT perform a synergistic action during systole: indeed, PM contract before LV wall contraction, allowing the stretching of CT, which elongate becoming taut [68]; this arrangement results in a better apposition of the MV leaflets and prevents prolapse as well as the inversion of MV cusps.

**Fig. 1.**
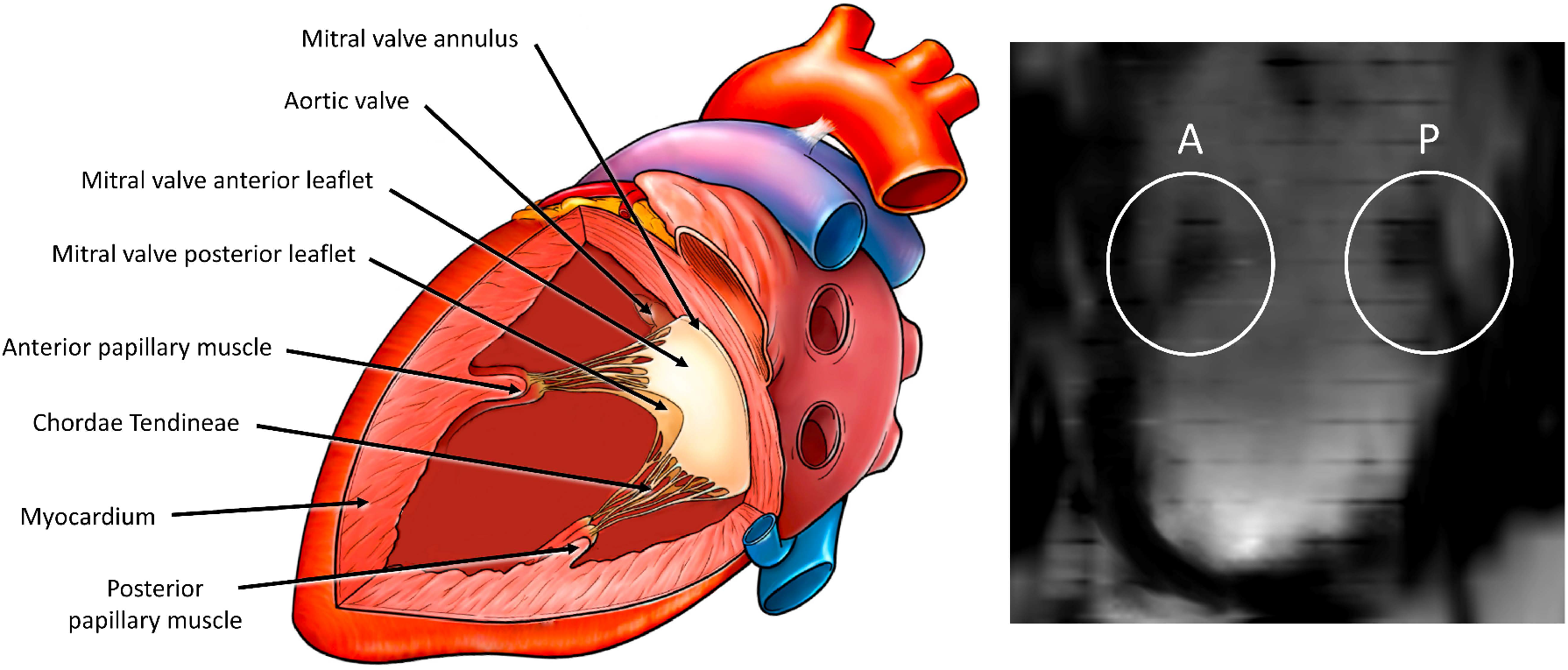
Left: Lateral view of the left ventricle, that emphasizes the anatomical structures of sub-valvular apparatus: papillaries protrude inside the ventricular cavity and they are connected to the two leaflets of mitral valve through chordae tendineae. Inspired by Nahabedian. Right: Cine-MRI vertical long axis through the left ventricle that shows the anterolateral (“A”) and the postero-medial (“P”) papillary muscles, which are structures arranged parallel to the longitudinal axis of LV [74].

In general, the effect of sub-valvular apparatus is ignored in the majority of cardiac hemodynamic computational studies; however, some recent works have considered the presence of papillary muscles, segmenting them from patient-specific magnetic resonance clinical images [19] or from high-resolution computer tomography [49, 92, 51]. Furthermore, few FSI studies have investigated the role of chordae models of different complexity, from a fibre-based elasticity modeling approaches (in which CT are modeled as thin elastic rods [54, 53, 36] or considering a discrete elastic pseudo-fiber model [37, 57, 59]) to fully 3d volumetric models (where CT are treated with an immersed boundary approach [33, 34, 26, 27, 28] or a smoothed particle hydrodynamic approach [88, 16, 17, 89]); in all these studies, CT are built using template geometries or segmenting them from patient-specific micro computer tomography, which is the only clinical imaging technique that has the sufficient spatial resolution to determine the CT length, the CT origins on the PM and the chordal insertion points on the MV leaflets [97].

The *mitral valve* is a bicuspid valve located between the left atrium (LA) and the left ventricle, characterized by an anterior and a posterior leaflets. Its role is to ensure the closure of the left atrioventricular orifice during systole and its proper opening to allow the ventricular filling in diastole. More in detail, during this latter phase, three different events can be recognized [63]: the *E-wave*, during which the valve reaches its maximum opening; the *diastasis*, characterized by a deceleration of the ventricular flow and partial valve closure; the *A-wave*, which corresponds to atrial systole and results in valve reopening. The succession of these three phenomena gives rise to the formation of complex vortex structures in LV that are responsible for a whole series of mechanisms triggering the LV remodeling [41, 67, 46].

In order to investigate the interaction of the ventricular blood flow with MV, two different mathematical modeling can be carried out. The first approach consists in using lumped parameter models to simulate valve dynamics, representing the fluid dynamics effect of the valve through the relationship between its pressure drop and flowrate [85, 48, 12, 66]; moreover, more sophisticated diode type models can be used, in which a time-dependent permeability is associated to the MV plane, in order to model the opening/closure of the valve without considering the leaflets [76, 77, 90, 21, 87]. The second approach consists in the 3D (or at least 2D) representation of the valve geometry, obtained either by parametric modeling [52, 22] or by segmentation of patient-specific real-time imaging data [95, 58, 96], whose presence can be considered through mesh conforming techniques (such as the Resistive Immersed Surface method [5, 78]) or fully Eulerian methods (such as the Immersed Boundary Method [69, 55, 34, 26, 27, 28] or the Fictitious Domain method [35, 56]). More in detail, this can be achieved by considering FSI between blood and valve leaflets, requiring the solution of a complex computational problem, where only the close (or open) valve configuration is required from imaging. Alternatively, one can consider DIB-CFD models also for the valve-fluid interaction [64, 31, 10], avoiding the complex FSI solution at the expense of the pre-processing of time-resolved valve imaging. In particular, the dynamics of the MV leaflets can be modelled in different ways: some studies employed an instantaneous opening/closure modality of the leaflets (i.e. *on/off*), neglecting the diastasis and the mitral reopening during the A-wave [18, 19, 9]; others, instead, prescribed an angle of opening and closure (taken from literature or by visual inspections of the medical images) simulating all the phases of diastole (E-wave, diastasis, A-wave and closure) [77, 84, 20] or only the opening (E-wave) and the closure [91, 20]. Finally, some authors reconstructed the MV in all the available frames of the medical images and prescribed the patient-specific displacement coming from all these configurations without the need of making assumptions on the opening and closure angles [8, 7].

In such contexts, in this work we provide a DIB-CFD study, based on time-resolved Cine-MRI images of a healthy and a MV regurgitant heart, with the aim of offering new insights in the modeling of sub-valvular apparatus and mitral valve dynamics. Specifically, the first aim of this work is to analyse the systolic hemodynamic role of the sub-valvular apparatus in a DIB-CFD framework. This allows us to provide useful information about the accuracy of the hemodynamics solution in presence of the complete sub-valvular apparatus. Indeed, at the best of our knowledge, the only works that consider CT in a DIB-CFD modeling [62] neglects the presence of PM. The second aim of this work is to compare the role of different mitral valve dynamics modelings: i) on/off modality; ii) prescription of physiological opening/closure time; iii) reconstruction of the mitral valve at four specific Cine-MRI image frames (peak of E-wave, peak of diastasis, peak of A-wave and fully closure). In particular, we are interested in determining whether scenarios i) and ii), in which the diastasis and the A-wave are neglected, can accurately replicate the physiological features and fluid dynamics patterns observed when diastasis and A-wave are included.

The outline of the work is as follows: in Section we briefly describe the reconstruction of the movement of the left heart (LH) internal wall starting from Cine-MRI imaging (Subsection 2.1). Furthermore, we present the mathematical framework (Subsection 2.2) and the numerical methods (Subsection 2.3), which will be exploited in Section 3 and Section 4 to computationally study the fluid dynamic role of sub-valvular apparatus and the different mitral valve dynamics.

## 2 Material and methods

### 2.1 Reconstruction of left heart geometries and motion from Cine-MRI imaging

This work studies, by means of Dynamic Image Based (DIB)-CFD, the fluid dynamics of patient-specific left hearts (LH), based on acquiring the endocardial cardiac motion through a preprocessing of dynamic medical images. To this aim, cardiac Cine-MRI data of two subjects are considered: a healthy subject (H) and a patient suffering from severe mitral valve regurgitation (R) due to a posterior leaflet prolapse. These data are provided by the Department of Radiology of Borgo Trento Hospital, Verona, Italy. Ethical review board approval and informed consent are obtained from all subjects.

More in detail, the image acquisition protocol and the reconstruction pipeline of left heart geometry and movement are described in our previous works [10, 9] and summarized in Figure 2. For each subject, the starting point is the elaboration of dynamic Cine-MRI images, consisting of 30 acquisitions per heartbeat (Figure 2A). Moreover, Figure 2 shows the two main innovations of the reconstruction pipeline, specific to this work: the segmentation and the motion reconstruction process of LV that allows to include also papillary muscles (Figure 2E, see Subsection 3.1) and the addition of chordae tendineae (Figure 2, see Subsection 3.2). The result is a time-discrete displacement field of the left heart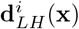, which is computed for each imaging frame (*i* = 1, …, 30) with respect to a reference configuration, i.e. the end systolic instant (Figure 2K). Since 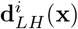is obtained only at the MRI acquisition times, a spline interpolation is employed to obtain a time-continuous displacement field **d**_*LH*_ (**x**, *t*).

**Fig. 2.**
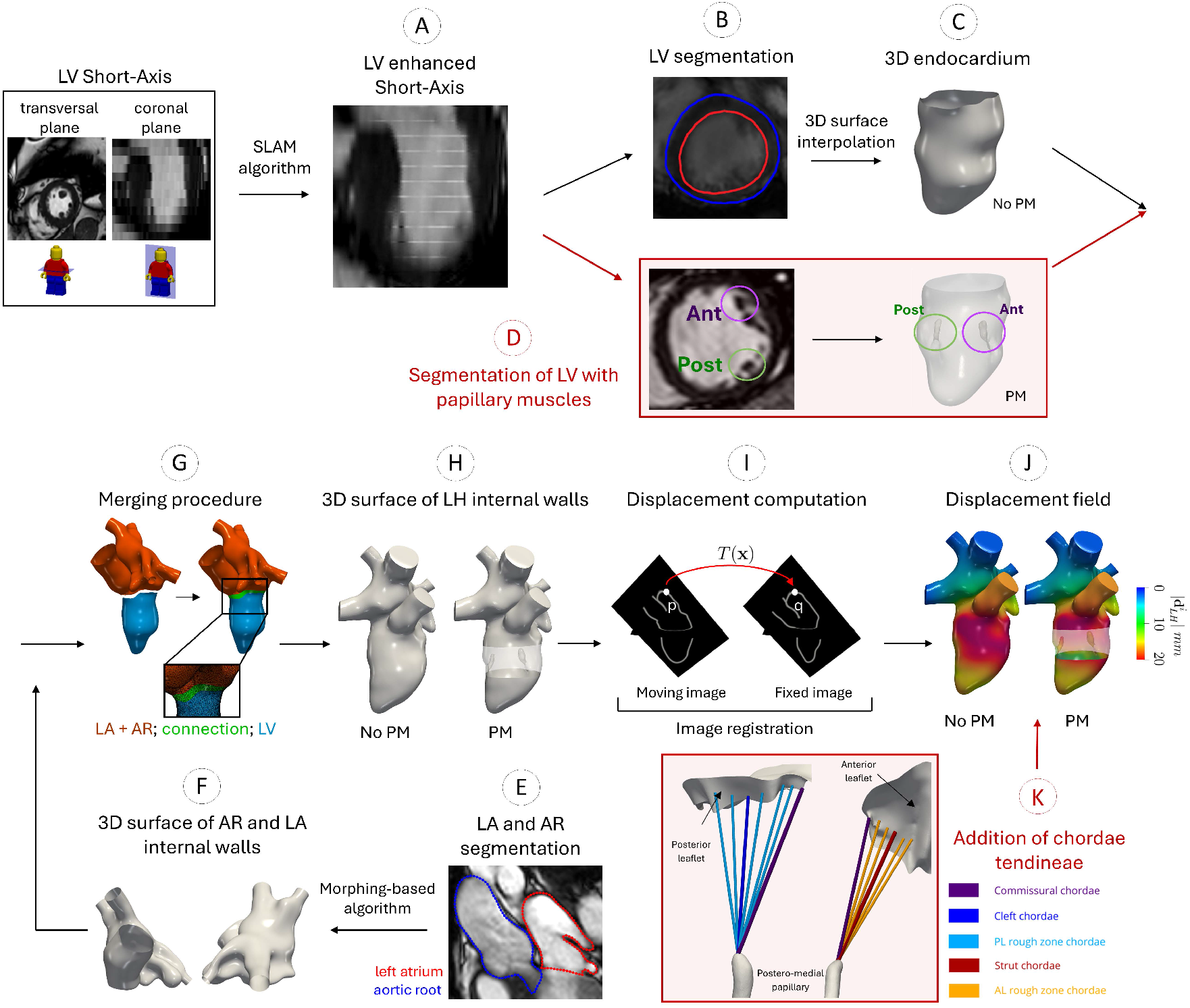
Schematic representation of the procedure employed to reconstruct the geometries and motion of the left heart from Cine-MRI imaging. A: Enhanced LV Short-Axis, obtained applying the Short-Long Axis Merging (SLAB) algorithm [32]. B: Segmentation of LV endocardium, manually detecting its contour (red). C: Endocardium surface obtained through a 3D surface interpolation [30]. D: Segmentation of both LV endocardium and papillary muscles (“Ant” refers to antero-lateral PM and “Post” to the postero-medial PM). E: Segmentation of aortic root and left atrium contour considering different Cine-MRI image series at disposal. F: 3D surfaces of AR and LA obtained through the morphing-based algorithm [75]. G-J: pipeline to obtain the displacement field for LH geometries: creation of a connection to merge the internal wall surface of LV with the lumen surfaces of AR and LA (G) and resulting 3D LH internal surfaces with and without PM (H); image registration procedure, in which a transformation operator T(x) maps a point p in a moving image to its corresponding point q within the fixed image (I); magnitude of LH internal wall surface displacement 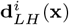 in the initial frame of systolic phase, with and without PM (J). K: Reconstruction of chordae tendineae, which are added to the geometries of reconstructed LH.

### 2.2 Modeling blood dynamics and ALE formulation

Blood is modeled as an incompressible, homogeneous and Newtonian fluid, described by the Navier-Stokes equations (NSE)[72, 71]. These equations have to be solved in a moving fluid domain, in order to account the heart contraction and relaxation during the cardiac cycle. Thus, the Arbitrary Lagrangian-Eulerian (ALE) framework [42, 23, 80] is considered; this formulation involves the introduction of a fictitious displacement field **d**_*ALE*_ associated to the fluid domain, obtained through a proper extension of the boundary movement **d**_*LH*_.

Inspired by [40, 3, 39, 15], **d**_*ALE*_ is obtained by extending **d**_*LH*_ into the fluid domain through the solution of a *non-linear lifting* problem, in which a quasi-incompressible Neo-Hookean fictitious material is associated to the mesh:

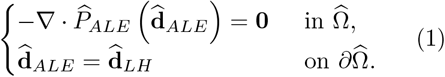

Here, 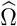 is the fluid domain reference configuration, and *P*_*ALE*_ is the first Piola-Kirchhoff tensor, defined as:

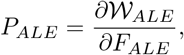

With

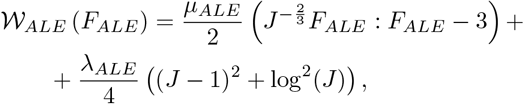

where *μ*_*ALE*_ and *λ*_*ALE*_ are the shear and the bulk modulus of the fictitious material associated to the mesh respectively, *F*_*ALE*_ = *∇***d**_*ALE*_ is the mesh deformation gradient, whose determinant is the jacobian *J*. Despite requiring a greater computational effort, the non-linear lifting ensures more robustness in the context of large deformation ALE problems than standard standard lifting operator (such as harmonic extension operator [101, 14] or linear elasticity [44, 80], even introducing local stiffening techniques [80, 43, 10, 9]), allowing to deform mesh elements without ruptures and irregularities.

At this point, it is possible to introduce the fluid domain velocity **u**_*ALE*_ and the ALE derivative operator *δ/δt*, defined through the Reynolds Transport Formula:

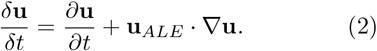

Denoting by Ω = Ω(*t*) the moving fluid domain, the strong formulation of the fluid dynamics problem for the domain Ω, shown in Figure 3, left, is: *Given the numerical parameters β, R*_*k*_, *ε*_*k*_, *K* = *{AV, MV, CT}, and the pressure functions p*_*in*_(*t*), *p*_*out*_(*t*), *for each t ∈* (0, *T*] *find blood velocity* **u**: Ω ∈ ℝ^3^ *and pressure p*: Ω ∈ ℝ *such that:*

**Fig. 3.**
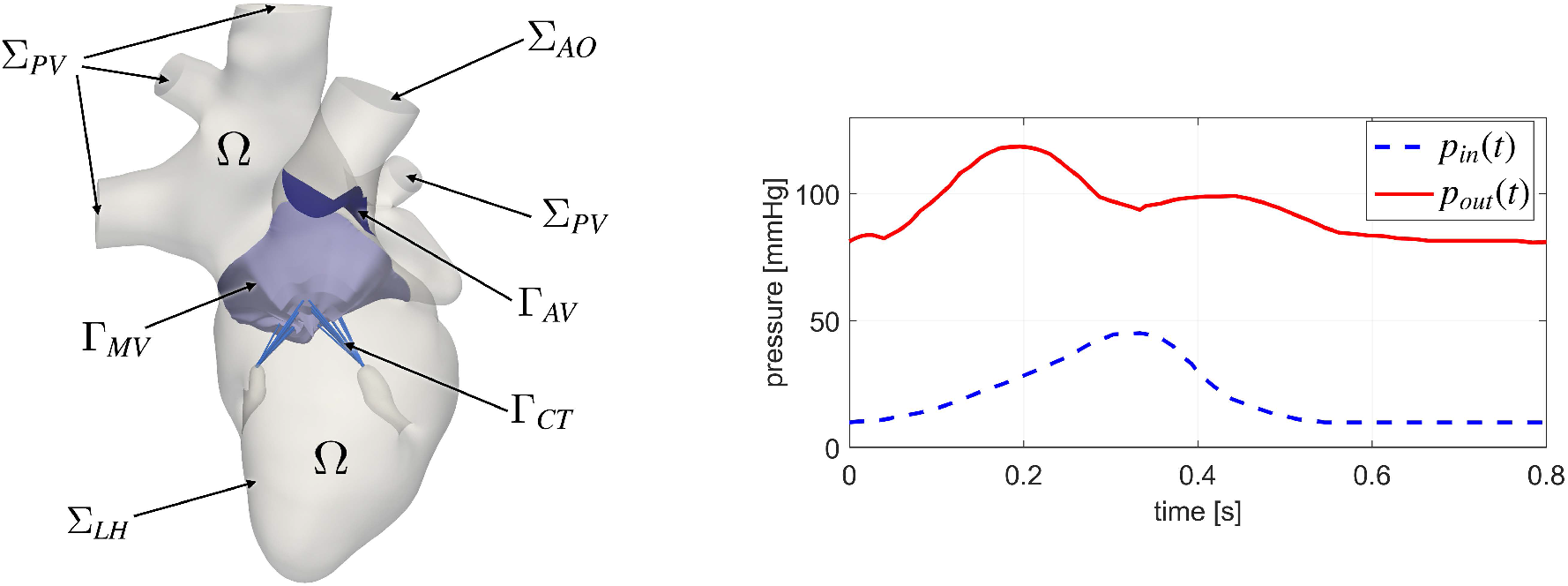
Left: Computational domain Ω with its boundaries and immersed surfaces. In light blue we report the mitral valve Γ_*MV*_, in dark blue the aortic valve Γ_*AV*_ and in blue the chordae tendineae Γ_*CT*_. Right: Pressures prescribed at the inlet Σ_*P V*_ and outlet Σ_*AR*_.

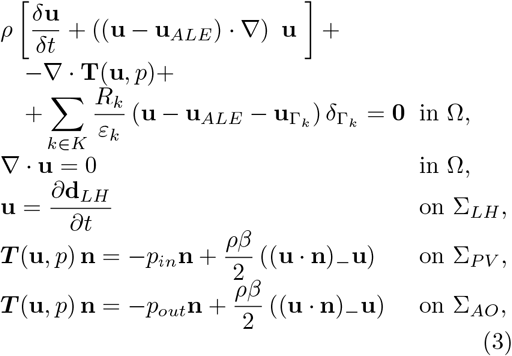

with null initial condition and where the boundary data *p*_*in*_(*t*), *p*_*out*_(*t*) are represented in Figure 3, right: in particular, the pressure prescribed on aortic root section Σ_*AO*_ is assumed to be the same for healthy and diseased cases and it is taken from the Wiggers diagram [99, 16]; whereas, at the pulmonary veins sections Σ_*P V*_ a constant pressure of 10 *mmHg* is imposed for healthy subject [99, 13], while a time-dependent pressure in case of diseased subject [16]. In particular, in the latter case, the formation of a regurgitant volume in the atrium during the systolic phase results in a marked increase of the atrial pressure [61].

In the above system, *Large Eddy Simulation (LES) σ-model* [64] is exploited to capture the transition to turbulence occurring in the left heart [18]. This implies that the Cauchy stress tensor for Newtonian incompressible fluid *σ*(**u**, *p*) is defined as:

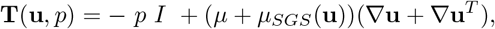

where *μ* is the dynamic viscosity and 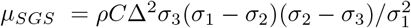 is the turbulent viscosity (sub-grid viscosity), with *ρ* the blood density, Δ the spatial filter w idth e qual t o the average mesh size, and *σ*_1_(**x**, *t*) *≥ σ*_2_(**x**, *t*) *≥ σ*_3_(**x**, *t*) the singular values of *∇****u***.

Chordae tendineae, mitral valve and aortic valve are treated as immersed surfaces (IS) Γ_*k*_ in the fluid d omain. T he i nteraction o f t he fluid with these surfaces is treated, by means of the last term in (3), with the *Resistive Immersed Implicit Surface* (RIIS) formulation [29, 25], which weakly enforces the blood to adhere to IS by means of a penalization technique, where *R*_*k*_ is the resistance parameter, *ε*_*k*_ is the half-thickness of the immersed surface, 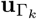 its velocity, while 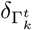 is a smoothed Dirac function, which implicitly describes the IS, defined as:

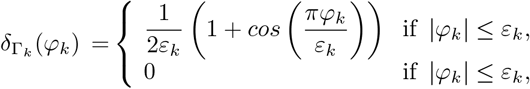

where *φ*_*k*_: Ω *→* ℝ is a level set function that describes Γ_*k*_ = *{***x** ∈ Ω: *φ*_*k*_(**x**) = 0*}*.

**Remark 1**. *Notice that in all the numerical experiments of this work we have made the quasistatic assumption to treat the resistive term in equation* (3). *This implies that* 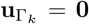*for k* = *AV, MV, CT*. *This means that we are accounting for the presence of the valve only from a geometric point of view (indeed, its encumbrance and position are included in the formulation by means of* 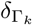 *term and they change in time). This choice has been suggested by stability issues arising for* 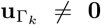 *and it is justified by noticing that it affects the fluid behavior only in a limited manner and only during the fast valve opening. For this reason, this choice has been widely used in previous works [25, 31, 32, 15, 10, 9]*.

Regarding the boundary conditions of Problem (3), we notice a no-slip velocity condition on Σ_*LH*_, while a backflow-stabilized Neumann condition[11] set on artificial sections, to prevent energy instabilities due to vortexes appearing at such boundaries.

### 2.3 Numerical methods and simulation setting

To numerically solve system (3), we employ a BDF first-order semi-implicit scheme [73] for time discretization and ℙ^1^ *−* ℙ^1^ Finite Elements for space discretization, together with SUPG/PSPG stabilization technique [86]. At each time step, the corresponding linear system is solved with a Simple preconditioned GM-RES method, implemented in the multi-physics high performance library *life*^*x*^ [1, 2] (https://lifex.gitlab.io/) based on the deal.II core [4].

Numerical experiments are run using 192 parallel processes on the GALILEO100 super-computer (https://www.hpc.cineca.it/hardware/galileo100) at the CINECA high-performance computing center (Italy).

We use the following values for physical and numerical parameters: *C* = 1.5, Δ*t* = 5 *·* 10^*−*4^ *s, ρ* = 1.06 *·* 10^3^ *kg/m*^3^, *μ* = 3.5 *·* 10^*−*3^ *Pa · s, μ*_*ALE*_ = 1.0 *·* 10^4^ *Pa* and *λ*_*ALE*_ = 1.0 *·* 10^2^ *Pa*. The latter value is relatively low and, consequently, it allows a certain deformation of mesh elements, but without distorting them excessively.

Tetrahedral meshes are built with dimension *h* determined through a mesh sensitivity analysis: an average mesh element size of 0.9 *mm* is obtained, with a refinement in the region of Left Ventricle Outflow Tract, LVOT (*h* = 0.75 *mm*) and papillaries (*h* = 0.55 *mm*). Furthermore, the RIIS treatment of immersed structures implies a local mesh refinement in the region of heart valves (*h* = 0.3 *mm*) and chordae tendineae (*h* = 0.2 *mm*). Moreover, we notice that the mesh resolution is able to satisfy the Pope criterion [70], i.e. we capture 80%, on average, of the turbulent kinetic energy.

## 3 Modeling the sub-valvular apparatus

This section is devoted to show the fluid dynamics role of the sub-valvular apparatus: in Subsection 3.1 the geometric segmentation and the motion reconstruction of papillaries starting from Cine-MRI patient-specific images is shown, while in Subsection 3.2 the design and inclusion of chordae tendineae is described. Finally, Subsection 3.3 reports the results of the numerical experiments, highlighting the main hemodynamic effects of sub-valvular apparatus during the systole, when CT and PM mainly interact with and influences blood dynamics. Notice that, for sake of simplicity, in the all sub-valvular apparatus simulations, the on-off dynamics of mitral and aortic valves is considered (see the D0 scenario in Section 4).

### 3.1 Geometric segmentation and motion reconstruction of the cardiac chamber including papillary muscles

In view of the reconstruction of the cardiac chamber for determining the fluid computational domain, this section aims to detail the papillary muscles segmentation and their motion reconstruction process, starting from patient-specific Cine-MRI images.

In particular, a segmentation procedure is designed to extract the left ventricle endocardium that accounts also the presence of PM, extending the left ventricle segmentation procedure proposed in Fumagalli et al. [31], which involves the use of a semi-manual segmentation algorithm [30], implemented in the Medical Image Toolkit (MITK) open-source software (www.mitk.org) [100, 65]. More in detail, the manual selection of endocardium has to take into account whether and how papillaries appear for each 2D transversal slice which composes the (time-resolved) 3D Cine-MRI images to segment. This implies three different ways to treat the image slices, depending on the position of PM (Figure 4A):

**Fig. 4.**
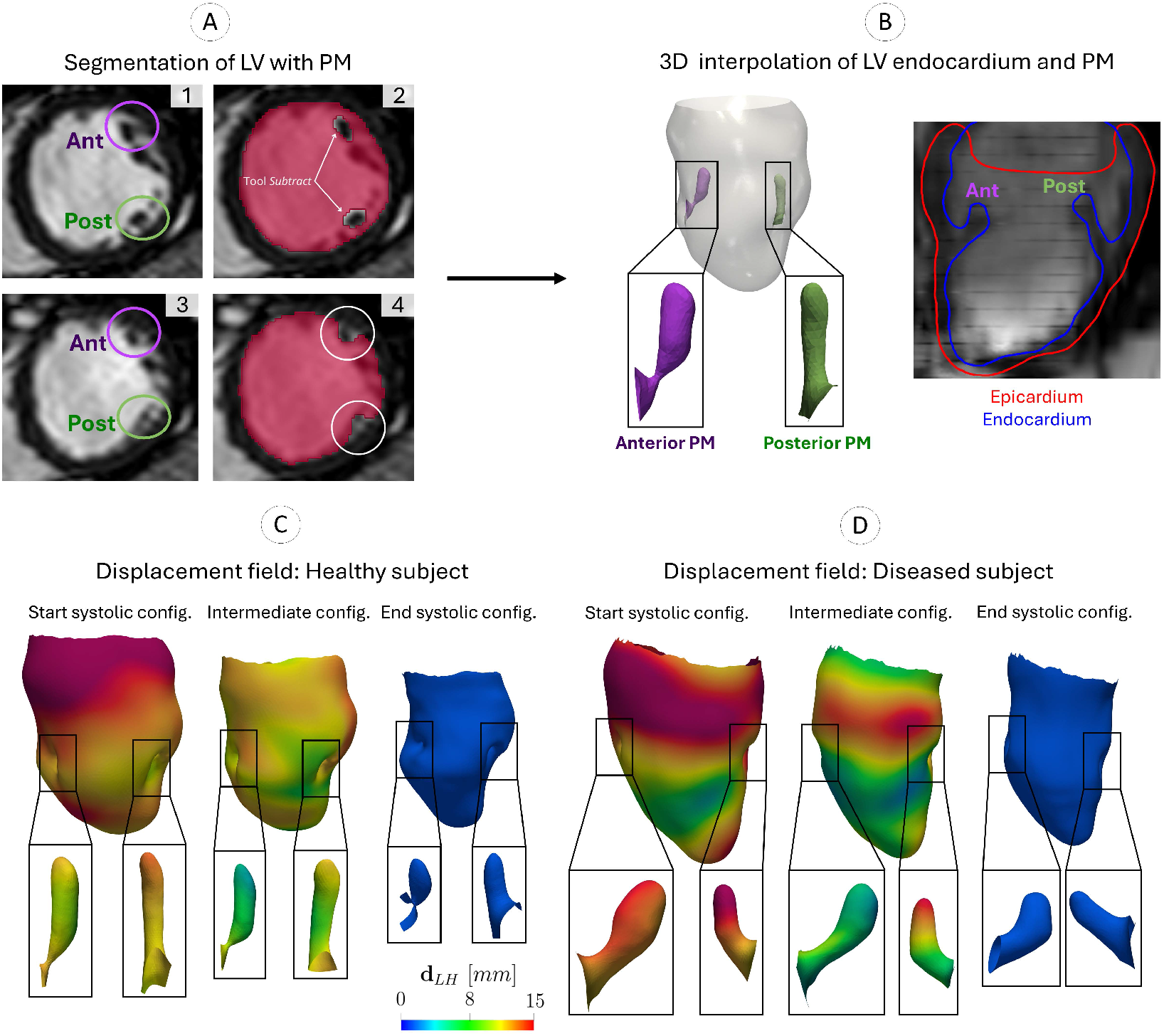
Schematic representation of the procedure employed to reconstruct the geometries of the cardiac chamber including papillary muscles (“Ant” indicates the antero-lateral PM, while “Post” the postero-medial PM). A: Segmentation of left ventricle endocardium accounting for PM reconstruction. In particular, in the 2D slices where papillaries appear inside the ventricular cavity (A1), their contours have to be excluded from the manual selection of the ventricular cavity (A2); while in slices where PM are attached to the cardiac wall (A3), the selection of endocardium considers also the PM contour (A4). B: 3D interpolation of the segmented endocardium accounting for PM reconstruction (left) and its visualization in a long axis view (right). C-D: Resulting displacement field **d**_*LH*_ of left ventricle endocardium that accounts for PM with respect to the end systolic configuration, at three representative frames, in case of healthy and diseased subjects. For each frame, a detail of the displacement of the antero-lateral (left) and postero-medial (right) PM external surface is shown.

1. In the first case, the image slice presents PM contours detached from the remaining endocardium (since they protrude inside ventricular cavity), thus papillaries look like two dark circles inside the white background of ventricular cavity (Figure 4A1). Then, once the region inside the endocardium, corresponding to the chamber, is identified (red area in Figure 4A2), PM are removed using the MITK *Subtract* tool, since they do not belong to the fluid domain;
2. In the second case, the image slice shows the PM attached to the cardiac wall; thus, a continuity between PM and endocardium contour is present (Figure 4A3). In this case, the selection of endocardium is done considering also the boundary of PM (Figure 4A4);
3. In the image slices where papillaries are not present, the internal region of endocardium is manually selected, identifying its contour.

Subsequently, the whole 3D area of the ventricular cavity is obtained by exploiting a radial basis function interpolation of the 2D contours, based on the image intensity [30] (Figure 4B). Thus, each segmented geometry presents the endocardium which accounts also the presence of PM. Afterwards, aortic root and left atrium are segmented using a multi-image based reconstruction algorithm [75] (Figure 2F-G) and merged to LV geometry (Figure 2H) in MeshMixer (https://www.meshmixer.com).

These steps are repeated for all the imaging acquisition frames. Then, we apply the motion reconstruction procedure proposed in [10, 9] (see Figure 2): the endocardium displacement of each frame is registered with respect to the end systolic configuration by exploiting the non-affine B-splines algorithm implemented in the Elastix open source library (http://elastix.isi.uu.nl) (Figure 2J-K) [47]. To perform an accurate registration, we apply it to the whole myocardium, obtained by segmenting also the epicardium (see Figure 4B).

The results of the reconstruction process are shown for three representative time instants in Figure 4 D-E, where the displacement field of endocardium (which accounts also papillaries) **d**_*LH*_ is shown at three representative systolic frames, in case of healthy and diseased subjects.

### 3.2 Design and inclusion of chordae tendineae

Since Cine-MRI imaging technique has not the sufficient spatial resolution that allows to visualize chordae tendineae, their external surfaces are reconstructed manually in the end diastolic configuration and then moved according to the mesh movement, participating to the ALE approach. More in detail, chordae tendineae surfaces are reconstructed as cylinders, departing from the tip of papillary muscles and attaching into different regions of the mitral valve. The quantity and the location of reconstructed chordae are chosen following the criteria of Lam’s functional classification [50]:

- 1 antero-lateral and 1 postero-medial *commissural chordae*: they insert respectively into the antero-lateral and postero-medial commissural areas of the mitral valve, i.e. the areas in which the two leaflets of MV are separated;
- 2 *cleft chordae*, that insert into the cleft zones of posterior leaflet, which are areas of free margin that divide the leaflet into three indentations;
- 9 anterior leaflet and 10 posterior leaflet *rough zone chordae*, which insert respectively in the rough zone of anterior leaflet and posterior leaflet of MV, i.e. a thick zone located adjacently to the leaflet free margin. Among these CT, 2 *strut chordae*, whose position evenly divide the rough zone of anterior leaflet, are also considered.

In these reconstructions, two important simplifications are considered: the absence of bifurcation, since the length and thickness of bifurcated chordae are not quantified in clinical and anatomical studies, and the absence of basal chordae, since both their number and their departing points from the left ventricle wall are not uniquely determined. Figure 5 shows a visualization of manually reconstructed chordae tendineae for the healthy and the diseased subjects. For the sake of simplicity, all chordae are built with a constant thickness, equal to 1 *mm*, regardless of chordae type. This allows us to use a single RIIS term to represent all chordae, with *ε*_*CT*_ = 0.5 *mm* and *R*_*CT*_ = 1.0*·*10^5^ *kg/m·s*. Further considerations on chordae thickness will be provided in Subsection 3.3.

**Fig. 5.**
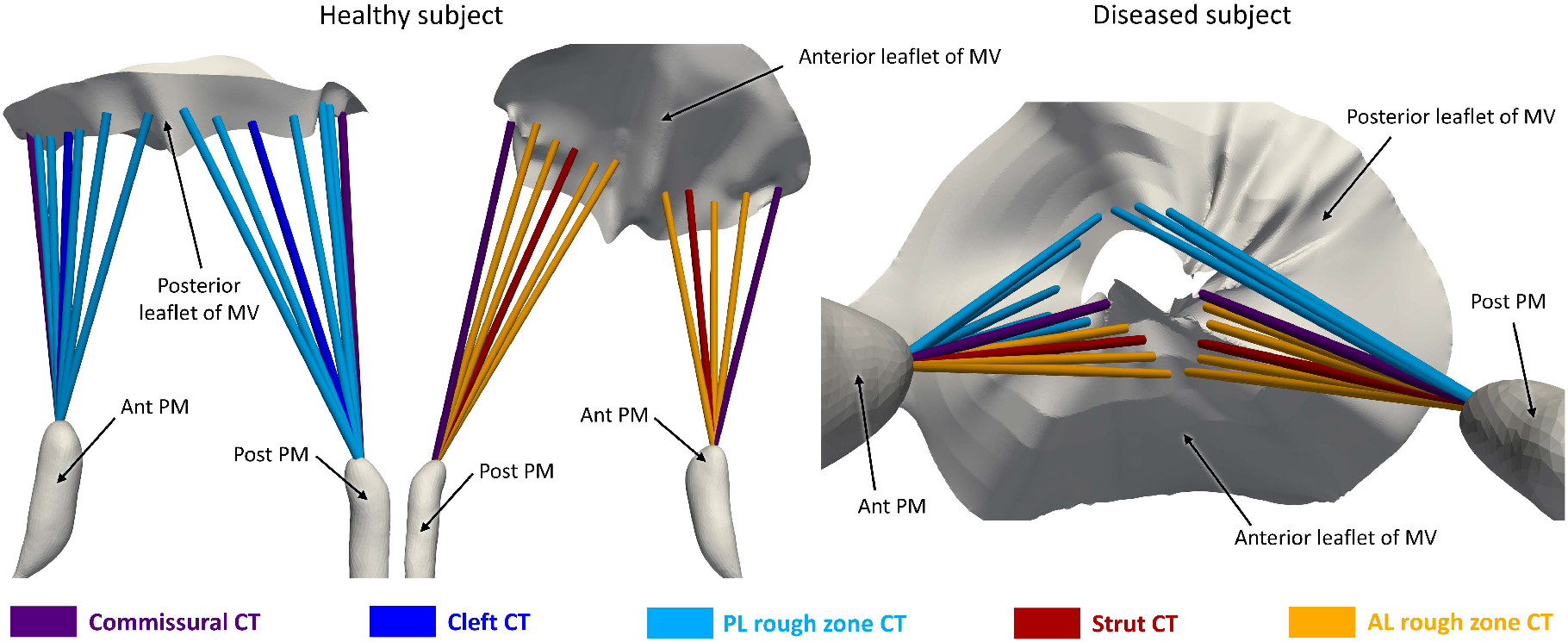
Global visualization of papillary muscles, mitral valve and manually reconstructed chordae tendineae, for healthy subject (left) and diseased patient (right), in the end diastolic configuration. According to the attachment position of CT in different region of MV, it is possible to distinguish among commissural, cleft, rough zone and strut chordae.

### 3.3 Numerical results of sub-valvular apparatus inclusion

This section shows the fluid dynamics results of the numerical experiments for the healthy subject (Subsection 3.3.1) and diseased patient (Sub-section 3.3.2), with the aim of determining the influence of chordae tendineae and papillary muscles on fluid dynamics (which will be discussed in Subsection 3.3.3). In particular, in order to discriminate the impact of these anatomical structures, three different scenarios are considered for each subject under study:

- *H_NoP_NoC* and *R_NoP_NoC* refer to healthy and diseased scenarios where the left ventricle endocardium is modeled neglecting the presence of papillary muscles and chordae tendineae. Thus, they are reference scenarios which represent the common configuration used in literature;
- *H_P_NoC* and *R_P_NoC* are healthy and diseased scenarios accounting for the presence of papillary muscles but excluding chordae tendineae;
- *H_P_C* and *R_P_C* refer to healthy and diseased scenario accounting for both the presence of papillary muscles and chordae tendineae.

We stress that this comparison is done during the systolic phase, when papillaries and chordae assume their maximum encumbrance, thus having a major role in influencing blood dynamics.

#### 3.3.1 Healthy subject

Firstly, the velocity field distribution is inspected, focusing on the local regions of papillary muscles and chordae tendineae.

Figure 6 shows the velocity streamlines at systolic peak (t=0.15 s) in the local region of papillaries, where the fluid particles locally surround them, in order to reach the LVOT region. More in detail, the presence (or absence) of chordae tendineae induces different velocity patterns in the region of PM: in a scenario without CT (Figure 6, H_P_NoC), the fluid streamlines pass over the tip of PM with a parallel direction to its surface, hitting it; instead, when CT are considered (Figure 6, H_P_C), the deflection of streamlines is much more consistent, especially in the region of PM tip. These different velocity patterns have an impact on the WSS acting on the surface of papillaries (Figure 6, right): since CT attach to the tip of PM, creating an obstacle with higher encumbrance, the blood tends to surround the PM-chordae complex, generating a lower WSS. Moreover, regardless of chordae presence, the antero-lateral PM surface exhibits a higher WSS with respect to posteromedial PM, since it is closer to LVOT region, therefore it is subjected to greater velocity gradients.

**Fig. 6.**
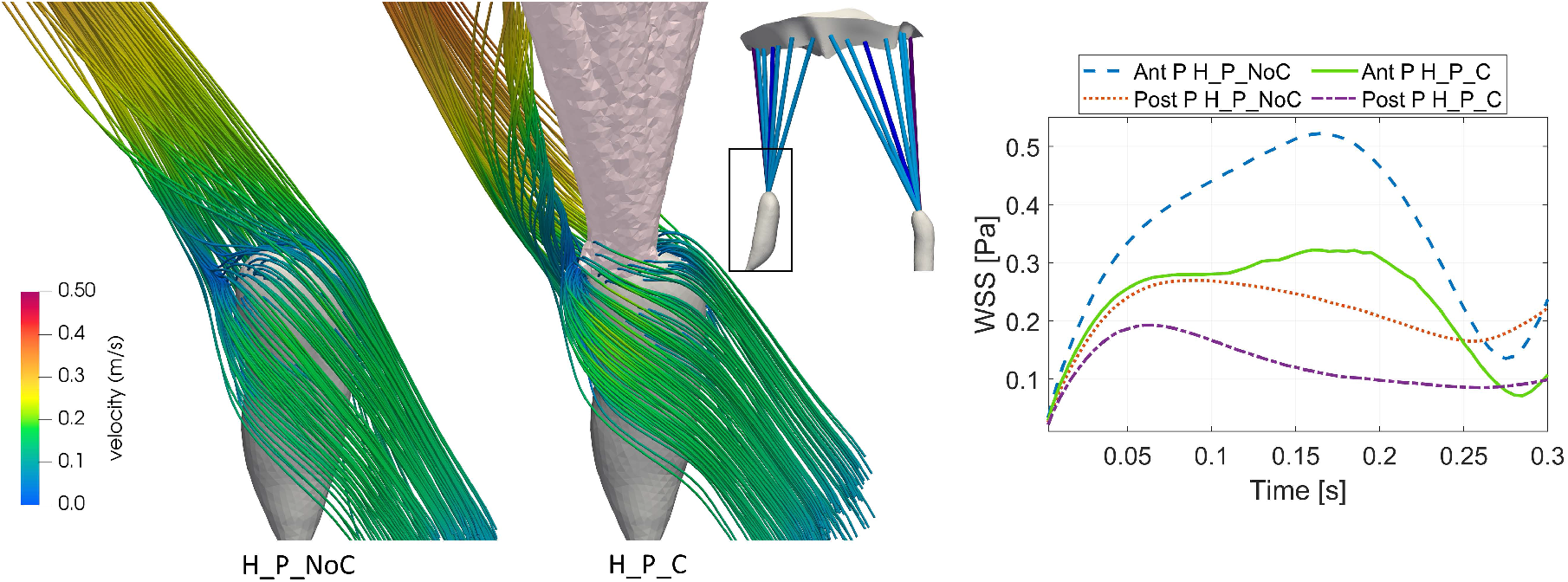
Left: Velocity streamlines in the region of anterior papillary muscle during the systolic peak for healthy case, in scenarios with papillaries and without (H_P_NoC) or with chordae tendineae (H_P_C). This region presents the chordae attachment to the tip of papillary muscles, thus they are grouped together creating a single structure (see the box). Right: Average wall shear stress acting on the surface of antero-lateral and postero-medial PM, during the entire systolic phase of the heartbeat. “Ant P” and “Post P” refer to antero-lateral and postero-medial PM in the two scenarios, respectively.

Chordae tendineae have an impactful effect on the velocity field variation also around the mitral valve. In particular, Figure 7A shows a top view of velocity streamlines as seen by the mitral valve, at systolic peak. In scenarios without CT (H_NoP_NoC and H_P_NoC, Figure 7A, left), the flow hits the entire surface of mitral valve, moving parallel to it in order to reach the LVOT. Instead, the presence of chordae reduces the quantity of flow that impinges the mitral valve, especially in the rough zone; this because the blood tends to surround the entire chordae assembly, rather than passing through the space among them (few streamlines with very low velocity magnitude are present in this region). Moreover, the local velocity pattern has a specific time evolution in the region of chordae which depart from anterolateral papillary muscle: from t=0 s to t = 0.12 s, after surrounding CT, streamlines flow in parallel, approaching the LVOT region; from t = 0.12 s until the end of systole, two vortexes, one clockwise and the other counterclockwise, are present (Figure 7A, center). These vortexes decrease in intensity and dimensions from mitral valve to papillaries region. This behavior does not concern flow in the zone of chordae which depart from posteromedial papillary muscle (Figure 7A, right), since parallel streamlines result for the entire systolic duration.

**Fig. 7.**
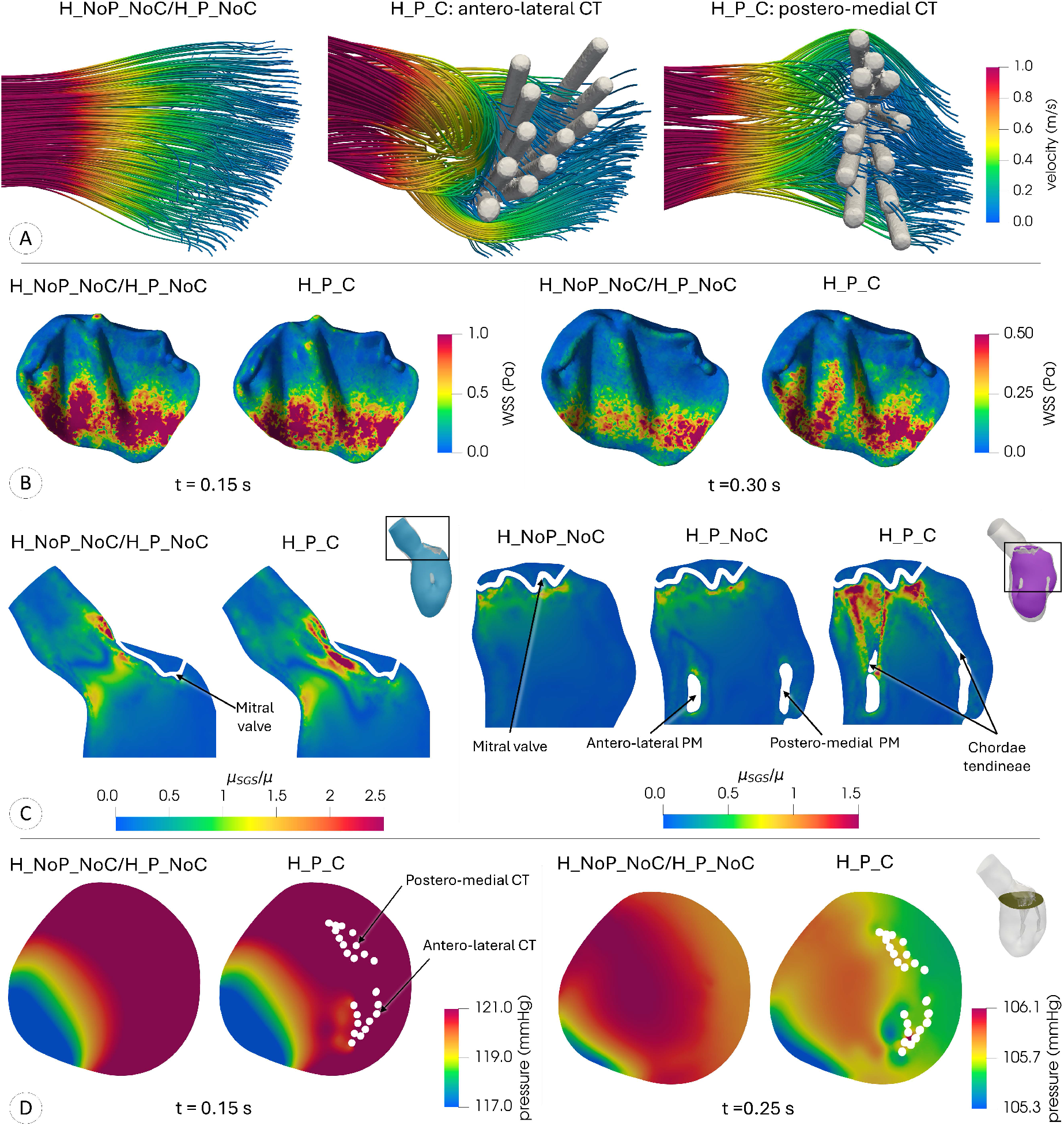
A: Local views of velocity streamlines, as seen by the mitral valve, at the instant of systolic peak. The scenarios without chordae (H_NoP_NoC and H_P_NoC scenarios) show a comparable velocity behavior, while the presence of chordae tendineae (H_P_C scenario) involve different velocity patterns. “H_P_C: antero-lateral CT” and “H_P_C: postero-medial CT” refers to chordae that depart from the antero-lateral PM and to the postero-medial PM, respectively. B: Spatial distribution of WSS field acting on the mitral valve, at the instant of systolic peak (t=0.15 s, left) and of the end systolic configuration (t=0.30 s, right). C: *μ*_SGS_*/μ* in LVOT region (left) and in a longitudinal section (right) at the instant of systolic peak for all the scenarios of healthy subject. D: Spatial distribution of pressure in a transversal section, at mitral valve level, at t=0.15 s (left) and at t=0.25 s (right). Legends obtained to highlight spatial variations within the no chordae scenario (H_NoP_NoC). Healthy case.

The previously described vortexes imply a different time behavior of WSS acting on the mitral valve among the scenarios which are considered. In particular, from t = 0 s to t = 0.205 s, the WSS is lower in chordae scenario, since CT are able to deviate the flow, ensuring lower velocity gradients in the region of their attachment to the valve (Figure 7B, left, t=0.15 s); instead, from t = 0.205 s until the end systolic configuration, the vortexes presence induces an increase of WSS on mitral valve, since their flow particles impinge it (Figure 7B, right, t=0.30 s). However, for all the scenarios, the maximum values of WSS occur in the region of anterior leaflet, particularly in the zone close to the LVOT, where the fluid particles undergo an increase of velocity.

The variations of velocity field due to the presence of papillaries and, especially, chordae tendineae imply also a different local development of turbulence, which is quantified through the turbulent viscosity *μ*_SGS_. Firstly, CT induce a different spatial distribution of this quantity in the LVOT region, where *μ*_SGS_ assumes its maximum values: in scenarios without CT the highest turbulence occurs in the aortic root region, while in chordae scenario it occurs in the area below the mitral valve (Figure 7C, left), since this is the region where streamlines, after surrounding CT, group together to reach the LVOT. Secondly, the presence of chordae implies a local increase of turbulence, that is shown on a longitudinal section at the instant of systolic peak (Figure 7C, right): in particular, high values of turbulent viscosity are reached in the region of chordae which depart from antero-lateral PM, where the two velocity vortexes develop. On the contrary, the presence of papillary muscles generates only a limited turbulent viscosity (less than half of dynamic viscosity).

Finally, the pressure spatial distribution is shown in a transversal section in proximity of mitral valve at two representative instants (Figure 7). In both the cases, we notice very small variations of the pressure field, which seems not so much influenced by the presence of the chordae. Similar arguments could be supported by the influence of the papillary presence on the pressure (the corresponding numerical results are not here reported for the sake of esposition).

#### 3.3.2 Regurgitant subject

The effect of chordae tendineae in reducing WSS acting on the mitral valve is even more evident in the case of mitral valve regurgitation.

Figure 8A shows velocity streamlines in the region of mitral valve at the instant of systolic peak (t=0.17 s). In the scenarios without chordae (R_NoP_NoC/R_P_NoC, Figure 8A, left), the velocity streamlines separate into an aortic jet (flowing toward the LVOT region) and a regurgitant jet (flowing toward the atrium due to mitral prolapse) in the region of mitral valve, hitting its surface. When the presence of chordae is considered (R_P_C, Figure 8A, right), the flow is split into two jets in a different way, in a region quite far from the mitral valve: the regurgitant jet evolves with vortexes and disturbance, entering inside the internal area created by the whole set of chordae; the aortic jet is able to surround the chordae at their middle length, avoiding to hit MV surface.

**Fig. 8.**
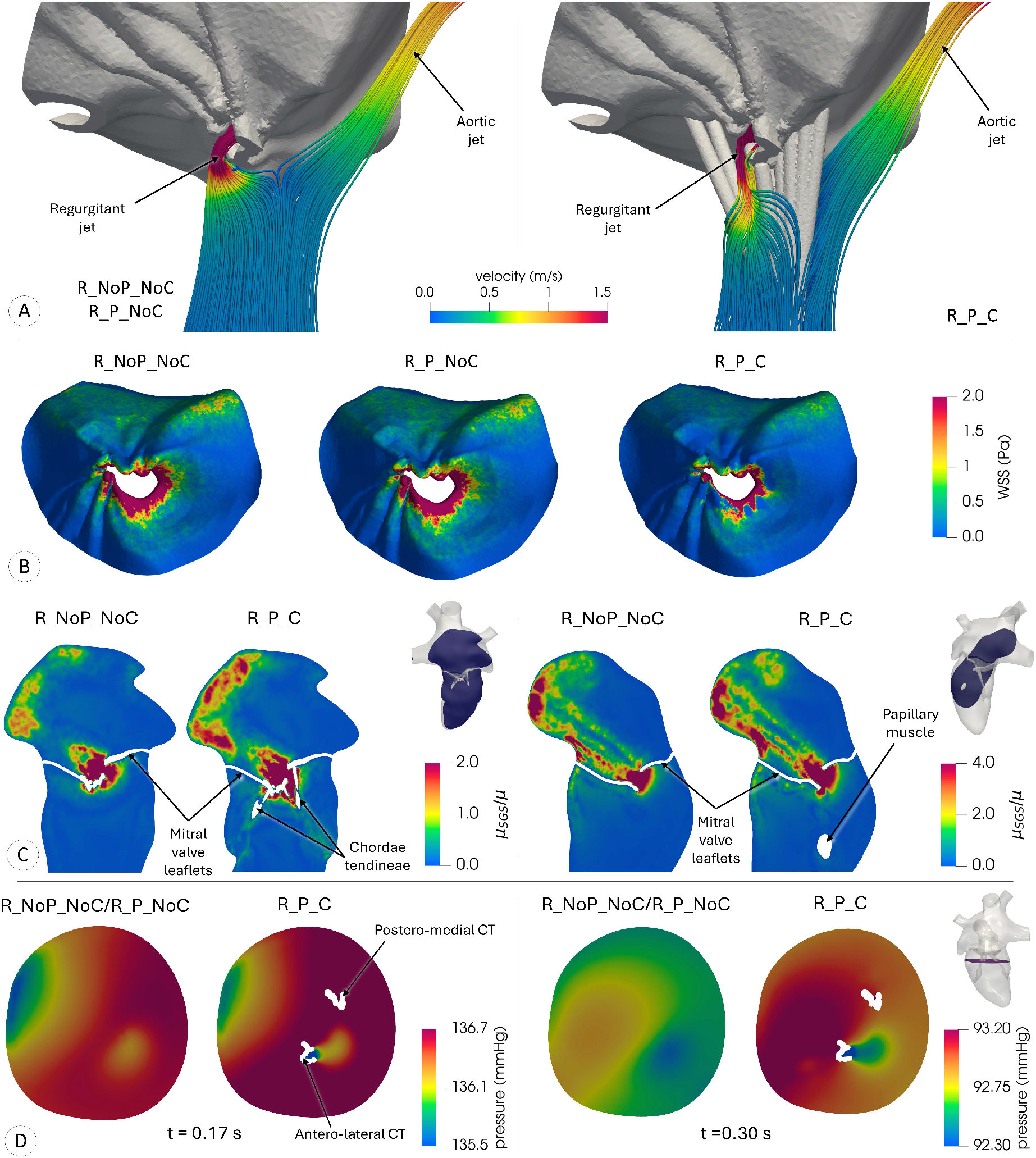
A: Velocity streamlines in the region of mitral valve and antero-lateral chordae tendineae at instant of systolic peak for the scenarios without chordae (R_NoP_NoC and R_P_NoC) and the scenario with chordae (D_P_CCost). B: Spatial distribution of WSS field acting on the mitral valve, at the instant of systolic peak, for all the scenarios of diseased patient. C: *μ*_SGS_*/μ* in a longitudinal section which includes chordae tendineae (left) and in the regurgitant section (right) at the instant of systolic peak. D: Spatial distribution of pressure in a transversal section, at mid-length chordae level, at t=0.17 s (left) and at t=0.30 s (right). Legends obtained to highlight spatial variations within the no chordae scenario (R_NoP_NoC). Regurgitant case.

These velocities patterns have consequences on WSS acting on mitral valve. In general, this latter present differences, in terms of entity and spatial distribution, with respect the healthy subject, since it is influenced by the presence of regurgitant jet and prolapse: since velocity assumes very high values in the region of atrioventricular orifice, the maximum WSS results in the MV area closer to free margin. More in detail, Figure 8B (right) shows the WSS distribution on MV at the systolic peak: in the scenario with chordae tendineae (R_P_C) there is a reduction in size and extension of the maximum WSS area compared to the scenarios without CT (R_NoP_NoC and R_P_NoC). This behavior is justified by the role that chordae tendineae play in diverting the flow: the absence of CT implies that the flow impinges directly the mitral valve, while CT presence represent a fluid dynamic obstacle which allows the flow to be divided away from MV; thus, a lower WSS affects its surface.

Mitral prolapse and regurgitant jet have also an impacting effect on turbulence. Indeed, considering a generic transversal section at the instant of systolic peak (Figure 8C, left), maximum values of *μ*_SGS_ are reached in the region of mitral valve prolapse, where turbulent viscosity exceeds dynamic viscosity more than 15 times. Moreover, this aspect is accentuated in chordae tendineae scenario (*R_P_C*): their presence implies more vortexes and disturbances which extend the turbulence in a wider region of space. In addition to this, turbulence develops also in the left atrium, due to the presence of regurgitant jet. Indeed, large turbulence is present in the region of regurgitant jet, not only in the region of prolapse, but also in the points where the jet impinges atrial wall (Figure 8C, right). Furthermore, *μ*_SGS_ assumes higher values at the atrium wall in chordae scenario with respect to scenarios without chordae.

Finally, also in case of diseased patient, spatial pressure variations due to chordae presence are very small (Figure 8D shows the pressure spatial distribution in a transversal section at the mid-length chordae level, at two representative instants). Similar arguments could be supported by the influence of papillary muscles presence on the pressure (the corresponding numerical results are omitted for the sake of esposition).

#### 3.3.3 Discussion on the inclusion of papillary muscles and chordae tendineae in cardiac hemodynamics

From the results of the previous sections, we notice a mutual interaction between the sub-valvular apparatus and blood dynamics. On one hand, CT and PM act as fluid dynamics obstacles, creating variations in local hemodynamic quantities (see, Figure 6, left, Figure 7A and Figure 8A). In particular, the chordae encumbrance is able to deviate the blood flow, generating peculiar velocity patterns which cause, in general, the reduction of WSS acting on MV (Figure 7B and Figure 8B). To better highlight this effect, we report the averaged-in-space WSS for both subjects in Figure 9, top. Furthermore, the chordae presence leads to an increase of turbulence, especially in mitral valve region, where CT have the highest encumbrance, causing disturbances and irregularities to the flow (see Figure 7C and Figure 8C). This increment is noticed also for the turbulence values averaged over the entire ventricle, reported in Figure 9, center.

**Fig. 9.**
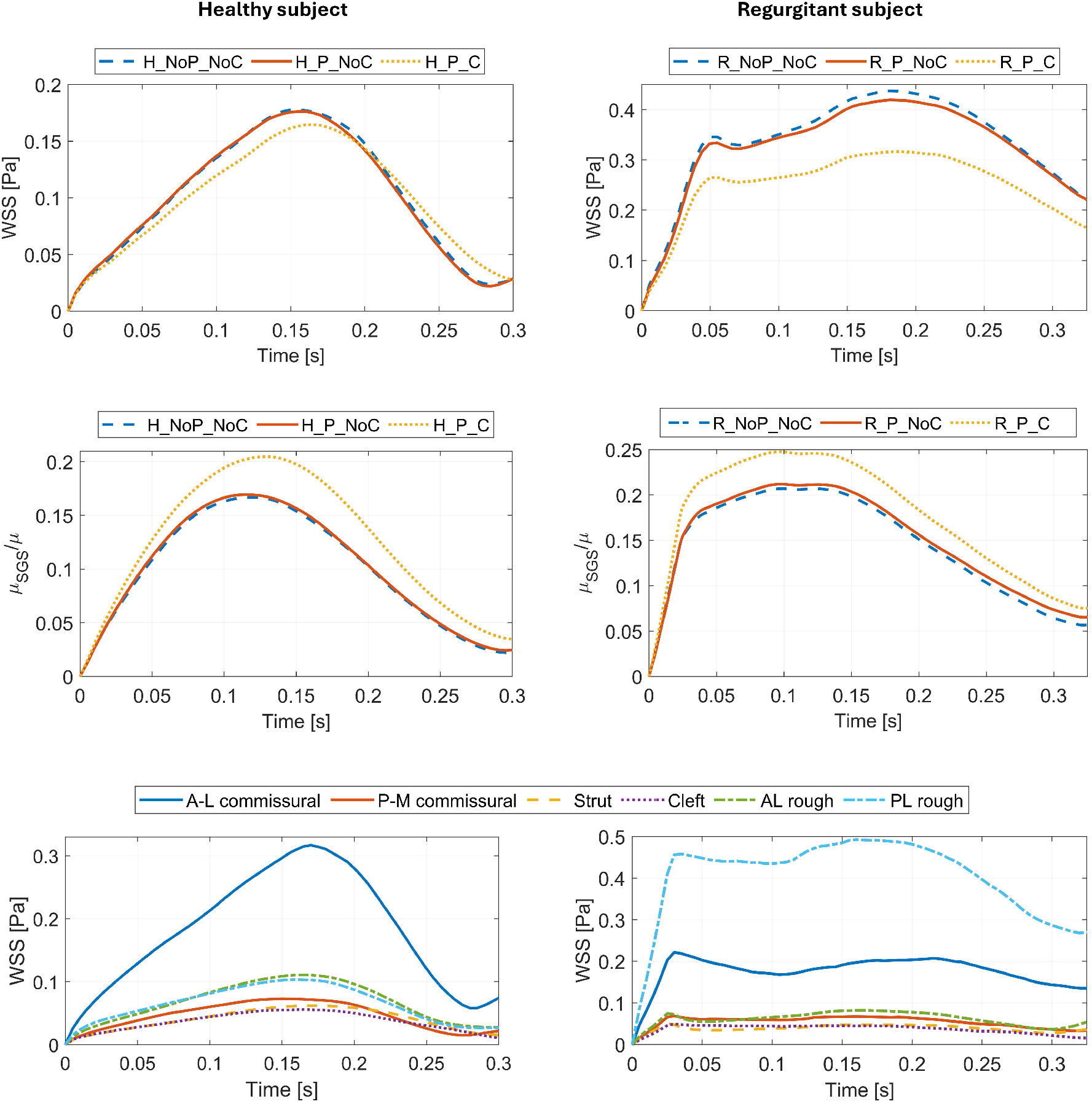
Top: Averaged-in-space WSS acting on the surface of mitral valve, during the systolic phase of heartbeat. Center: Time behavior of normalized turbulent viscosity, averaged in the whole ventricle. Bottom: Average WSS acting on the surface of chordae tendineae, distinguished depending on the type of chordae, whose distinction is described in Subsection 3.2. Left: healthy subject. Right: regurgitant subject.

On the other hand, it is important to assess also the effect of blood dynamics on papillaries and chordae. For example, blood dynamics exerts forces on the sub-valvular surfaces, see the spatial averaged WSS acting on chordae tendineae, reported for each type of chorda in Figure 9, bottom. Its behavior differs according the subject under study. In the healthy case, maximum values are reached at the instant of systolic peak, when velocity is particularly high (Figure 9, bottom-left); specifically, the antero-lateral commissural chorda is subjected to the highest WSS, since it undergoes the pronounced velocity gradients of the LVOT region and the effects of clockwise and counterclockwise velocity vortexes, described in Subsection 3.3.1. Instead, in case of the diseased patient, the average WSS has two peaks (the early one relates to the developing of regurgitant jet, while the late one to the systolic peak) and assumes higher values than healthy case, due to the elevated velocities that occur in the region of the prolapse (Figure 9, bottom-right). In this case, the highest WSS acts on the surfaces of posterior-leaflet rough zone chordae, since these CT undergo a elongation due to the MV prolapse, which causes an increase of their surface in contact with the fluid. Moreover, they are the most affected by the vortexes and fluid dynamic disturbances which develop due to the regurgitant jet.

Nevertheless, the inclusion of chordae tendineae and papillary muscles into DIB-CFD models seems to be less relevant if one is interested on global fluid dynamics quantities (the corresponding numerical results are not here reported for the sake of esposition). Furthermore, the pressure distribution is very similar with and without the CT and PM presence, and the pressure drops due to these structures are negligible for the entire systolic duration (see Figure 7D and Figure 8D).

Summarizing, we can state that the inclusion of papillary muscles and chordae tendineae into DIB-CFD models is recommended in case of specific investigations of the local fluid dynamics close to the sub-valvular apparatus.

Finally, we observe that we also considered the case of variable chordae thickness to account for the more physiological scenario. The results were found to be very similar to the “homogeneous” case showed in this paper, thus they are not here reported.

## 4 Modeling mitral valve dynamics

This section aims at analyzing the effect of different mitral valve dynamics models in the context of DIB-CFD framework. We observe that, in any case, the valve dynamics is not modeled by mean of a differential problem; rather the position of the leaflets is suitably obtained by Cine-MRI images.

### 4.1 Description of models and scenarios

Four valve dynamics modeling approaches are considered (refer for the representative healthy case to Figure 10A):

**Fig. 10.**
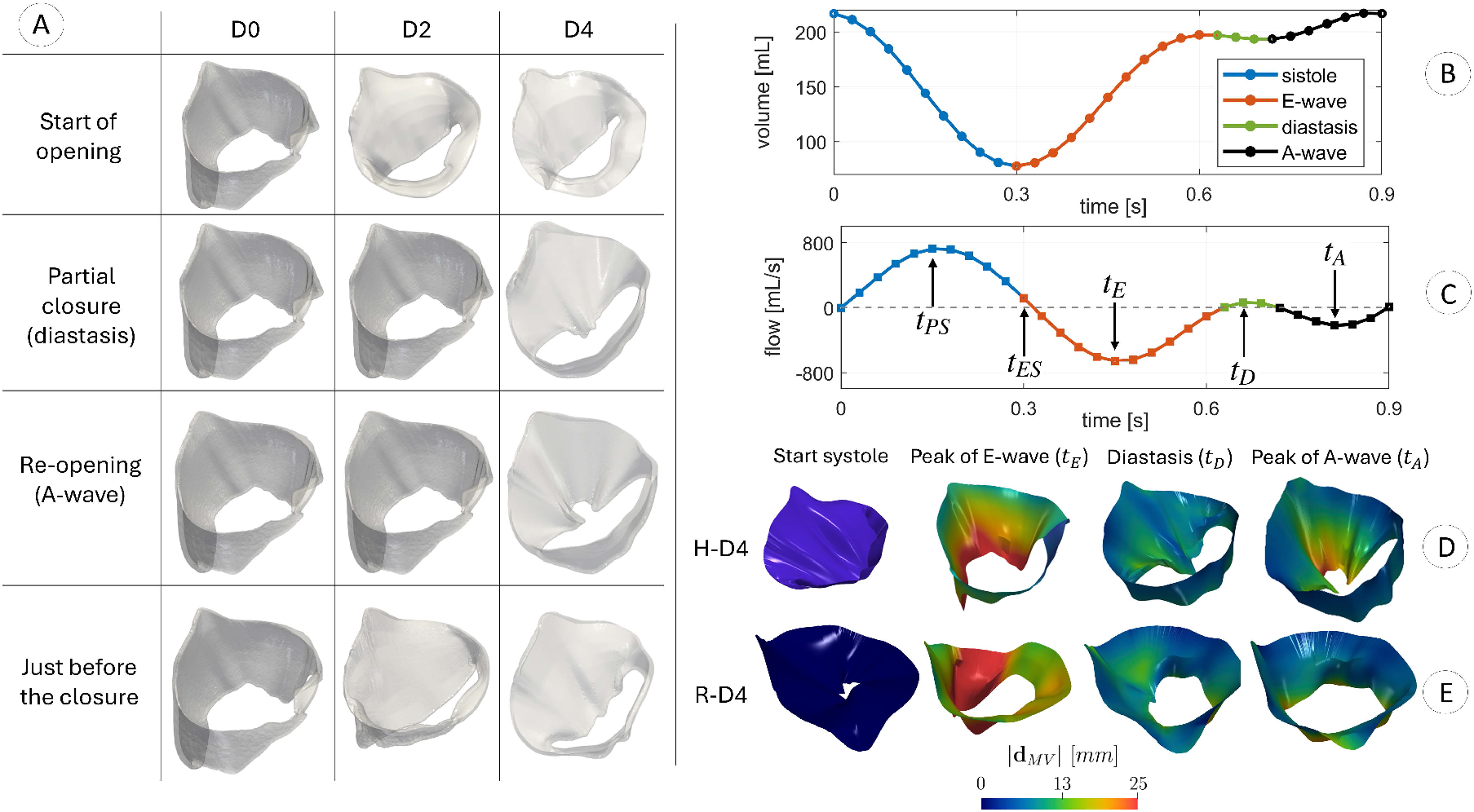
A: Mitral valve configurations for the three scenarios for the healthy subject at selected time instants of the diastolic phase. B: Variation in time of the reconstructed ventricular volume in case of healthy subject, see Subsection 2.1. C: Corresponding flow rate through the aortic (from 0 to 0.3 s) and mitral (from 0.3 to 0.9 s) orifices in case of healthy subject. The arrows refer to the instants of peak systole (*t*_*P S*_), end systole (*t*_*ES*_), peak of E-wave (*t*_*E*_), peak of diastasis (*t*_*D*_) and peak of A-wave (*t*_*A*_). D-E: Magnitude of the displacement **d**_*MV*_ obtained by imaging at four representative temporal instants, computed with respect to the systolic configuration. Scenario D4 for the healthy (D4_H) and regurgitant (D4_R) subjects.

- *D0 model* (*on-off modality*): instantaneous opening and closure of the valve, completely neglecting its dynamic. This implies that only two valve configurations (the fully open and the fully closed ones) are reconstructed;
- *D2 model*: the opening and closure dynamic is accounted by using a time interpolation of the fully open and fully closed configurations on a selected time interval;
- *D4 model* (only for MV): four specific imaging frames are considered and then suitably time interpolated (shown in Figure 10B-C): the fully open and the fully closed configurations (as for D0 and D2), the peak of diastasis (MV partial closure), and the peak of A-wave (MV re-opening);
- *D30 model* (only for MV and healthy case): all the available imaging frames are considered and then suitably time interpolated.

We observe that the D30 model, when available (i.e. for the MV healthy case), is used as *gold standard* to assess the accuracy of D0, D2, D4 MV scenarios.

For each model, MV geometries are segmented for the configuration of interest from dynamic rotated Cine-MRI images, acquired applying the protocol reported in [82]. Specifically, for each frame of interest the MV geometry is derived by employing the method described in [82, 10], based on tracing the valve leaflets on several planes and allowing to obtain a surface mesh of triangles.

We notice that for D0 and D2 models the position of the mitral annulus is in accordance with the ALE displacement, whereas for D4 and D30 by a suitable interpolation of the images at disposal. For the aortic valve, in this work we consider only D0 and D2 models, since we have a disposal only the opening and closing configurations of this valve from Cine-MRI images (we don’t have rotated images for the aortic valve).

Regarding the opening and closure time intervals, for D2, D4 and D30 models they are obtained by inspecting the Cine-MRI images, refer to Table 1. Instead, for the D0 models they are set equal to the time discretization step Δ*t* used in numerical experiments, since the valves dynamics is completely neglected, implying instantaneous opening and closure time intervals.

**Table 1.** Description of the scenarios as a function of AV and MV model, together with the corresponding time intervals of opening Δ_*op*_ and closure Δ_*cl*_ of aortic and mitral valves in healthy and regurgitant cases, for all the scenarios under study. In case of H-D0 and R-D0 scenarios, these intervals are set equal to the time discretization step Δ*t* used in numerical experiments (according to the definition of D0 model), while in the other scenarios they are obtained by inspecting the Cine-MRI images.

| Scenario | Model |  | Mitral valve |  | Aortic valve |  |
| --- | --- | --- | --- | --- | --- | --- |
| | MV Model | AV Model | $\Delta_{op}$ [ms] | $\Delta_{cl}$ [ms] | $\Delta_{op}$ [ms] | $\Delta_{cl}$ [ms] |
| H-D0 | D0 | D0 | $\Delta t$ | $\Delta t$ | $\Delta t$ | $\Delta t$ |
| H-D2 | D2 | D2 | 60 | 60 | 19 | 47 |
| H-D4 | D4 | D2 | 60 | 60 | 19 | 47 |
| H-D30 | D30 | D2 | 60 | 60 | 19 | 47 |
| R-D0 | D0 | D0 | $\Delta t$ | $\Delta t$ | $\Delta t$ | $\Delta t$ |
| R-D2 | D2 | D2 | 90 | 60 | 19 | 47 |
| R-D4 | D4 | D2 | 90 | 60 | 19 | 47 |

Starting from the geometric configurations at disposal, and applying the proper opening/closure time intervals, for D2, D4 and D30 models we perform a spline-time interpolation in order to describe the opening and closure dynamics. In Figures 10D-E, see also as an example the magnitude of the MV displacement for model D4 for both subjects under study.

In this work we considered the following scenarios (refer also to Table 1):

- D0 for MV and D0 for AV, for both the healthy and the regurgitant cases (in what follows referred to as *H-D0* and *R-D0*, respectively);
- D2 for MV and D2 for AV, for both the healthy and the regurgitant cases (in what follows referred to as *H-D2* and *R-D2*, respectively);
- D4 for MV and D2 for AV, for both the healthy and the regurgitant cases (in what follows referred to as *H-D4* and *R-D4*, respectively);
- D30 for MV and D2 for AV, for the healthy case (in what follows referred to as *H-D30*).

Both the valvular surfaces are treated with RIIS approach (Eq. 3), with *R*_*AV,MV*_ = 1.0 *·* 10^5^ *kg/m · s* and *ε*_*AV,MV*_ = 0.75 *mm*.

Notice that, for sake of simplicity, the following comparisons are based on numerical experiments where we do not include the sub-valvular apparatus.

### 4.2 Numerical results of the different mitral valve dynamics

This section shows a comparison of the numerical results obtained considering different scenarios introduced in the previous subsection: Subsection 4.2.1 and Subsection 4.2.2 show the results of healthy and diseased subjects respectively, while Subsection 4.2.3 presents a brief discussion of the results highlighting the importance of MV model in the numerical description.

To make this comparison, the following variables are introduced:

- *Ensemble* velocity and pressure, i.e. their average computed over 5 heartbeats;
- *Turbulent Kinetic Energy* (TKE), defined as the square root of the standard deviation of the ensemble velocity. It allows to quantify and localize the regions characterized by marked transition to turbulence [93, 81];
- *Global Turbulent Kinetic Energy* (GTKE), i.e. TKE integrated over the left ventricle. It allows to identify the instants where (in average) high velocity fluctuations occur by means of the fluid Reynolds stress tensor [19];
- *Time Averaged WSS* (TAWSS) and *Relative Residence Time* (RRT). This latter is a function of space providing a surrogate information about the effective residence time spent by particles close to the myocardial wall [38].

#### 4.2.1 Healthy subject

This subsection describes the comparison results among the four different scenarios (H-D0, H-D2, H-D4 and H-D30, refer to Table 1) in case of healthy subject.

Figure 11 (left) shows the magnitude of the ensemble velocity computed among five heart-beats and evaluated at four time instants of the heartbeat. We observe that at the systolic peak *t*_*P S*_ the velocities in LVOT region are comparable in the four cases, but their patterns seem to be slightly more chaotic in H-D4 and H-D30 scenarios. At the peak of E-wave *t*_*E*_ all the four scenarios feature two vortexes below the mitral valve leaflets, but different velocity patterns characterize H-D0 scenario, which does not experience the formation of a vortex in the middle of the ventricle. Moreover, H-D4 and H-D30 scenarios exhibit higher velocity in the local region of MV during the entire duration of the E-wave, as reported in Figure 12, top-left. At the peak of diastasis *t*_*D*_ the mitral valve undergoes a partial closure in H-D4 and H-D30 scenarios, whereas it remains fully open in the remaining two cases. This affects the velocity patterns in the ventricle: specifically, in H-D4 and H-D30 scenarios a uniform clockwise vortex develops in the central region of ventricular cavity; instead, in H-D0 and H-D2 the mitral valve is not able to address the flow towards the left ventricle wall, promoting a different vortexes dynamics. At the peak of A-wave *t*_*A*_ the second injection of fluid in the ventricle results much more consistent in D4 and D30 scenarios, which present higher velocity magnitude and comparable fluid dynamics patterns.

**Fig. 11.**
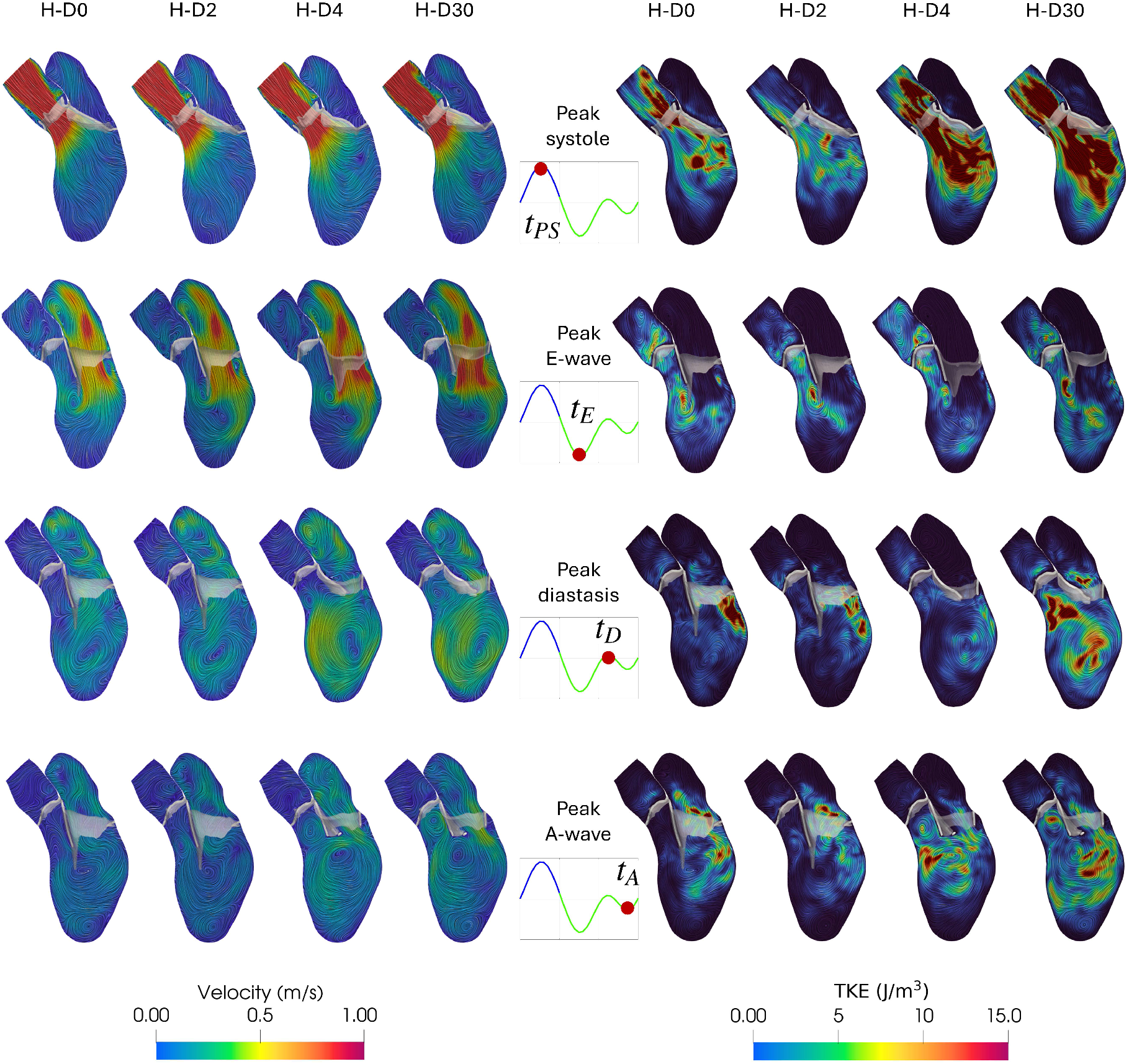
Magnitude of the ensemble velocity field (left) and spatial distribution of TKE (right) computed over five heartbeats at the peak of systole (*t*_*P S*_), peak of E-wave (*t*_*E*_), peak of diastasis (*t*_*D*_) and peak of A-wave (*t*_*A*_) in the four scenarios. The flow rate diagrams refer to the flow through the aortic (from 0 to 0.3 s, in blue) and the mitral orifice (from 0.3 to 0.9 s, in green). Healthy subject.

**Fig. 12.**
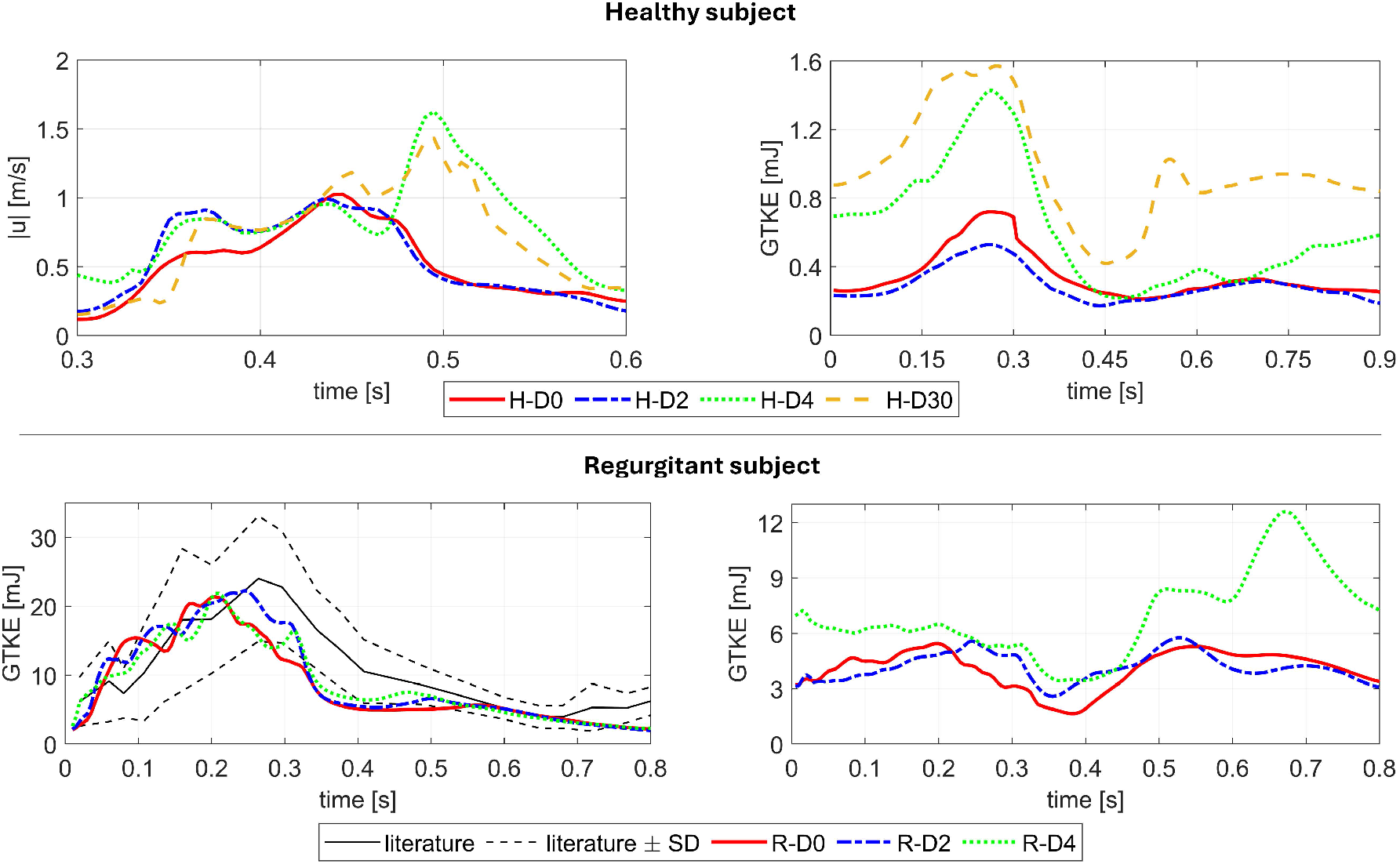
Top-left: Ensemble velocity magnitude in the region of the mitral valve orifice, computed in the entire duration of the E-wave for all the scenarios of healthy subject.Top-right: Time behavior of GTKE integrated over the left ventricle volume during the heartbeat for each scenario of healthy patient. Bottom: Evolution in time of GTKE integrated over the atrial (left) and ventricular (right) volume during the heartbeat in the three scenarios performed on diseased patient. For the left figure, a comparison with respect average values among several patients with mitral valve regurgitation by means of phase-contrast MRI [24] is reported.

Figure 11 (right) shows the spatial distribution of TKE computed throughout five heartbeats at the same temporal instants. At *t*_*E*_ all the scenarios experience a similar TKE pattern with large values below the anterior leaflet, exactly in correspondence of velocity vortexes detected in Figure 11 (left). At *t*_*D*_ H-D30 scenario features larger velocity fluctuations than the other three cases, especially in the middle of the ventricle and also in correspondence of LVOT. Conversely, H-D0 and H-D2 scenarios experience their maximum values below the posterior leaflet, while H-D4 scenario has the lowest TKE values among all scenarios. At *t*_*A*_ the maximum turbulence occurs in the middle of the ventricle in H-D4 and H-D30 scenarios, and close to the mitral annulus and along the ventricle wall in H-D0 and H-D2. Moreover, we observe that for H-D4 and H-D30 the amount of turbulence is increased due to the effect of modeling the diastasis, i.e. the partial closure of the valve, which creates a reduction of the area of the exiting mitral jet. At *t*_*P S*_ H-D4 and H-D30 scenarios feature a larger amount of turbulence with respect H-D0 and H-D2 scenarios. Probably, this is an effect of the increased amount of turbulence due to the valve closure at the diastasis, which propagates throughout the heartbeats.

Observing the GTKE time evolution in Figure 12, top-right, we found that regardless of MV dynamic modeling, the largest amount of turbulence occurs during the systolic phase of heartbeat. Moreover, higher values of GTKE results in H-D4 and H-D30 scenarios and this reflects the higher values of TKE at the peak of systole (Figure 11, right). During the diastolic phase, H-D30 features always the largest values, with its peak during diastasis (around 0.56 s). Conversely, the other three scenarios show comparable values during the E-wave and early diastasis. Nevertheless, scenario H-D4 experiences an increase of turbulence than H-D0 and H-D2 scenarios, towards the end of the cardiac cycle, as also showed for TKE in Figure 11 (right) at the peak of A-wave.

Figure 13 (top) shows the spatial distribution of the ensemble TAWSS and RRT. Their pattern in left atrium are quite similar among the four scenarios. In contrast, considering left ventricle, H-D4 and H-D30 scenarios exhibit high TAWSS values along the ventricular wall, attributed to the pronounced vortexes formed during diastasis (refer to Figure 11). This implies that, since the shear forces are able to washout possible stagnant flow, a reduction of RRT results in H-D4 and H-D30 scenarios with respect H-D0 and H-D4, suggesting that the inclusion of diastasis and A-wave could better describe the washout process developed by the left ventricle during diastole.

**Fig. 13.**
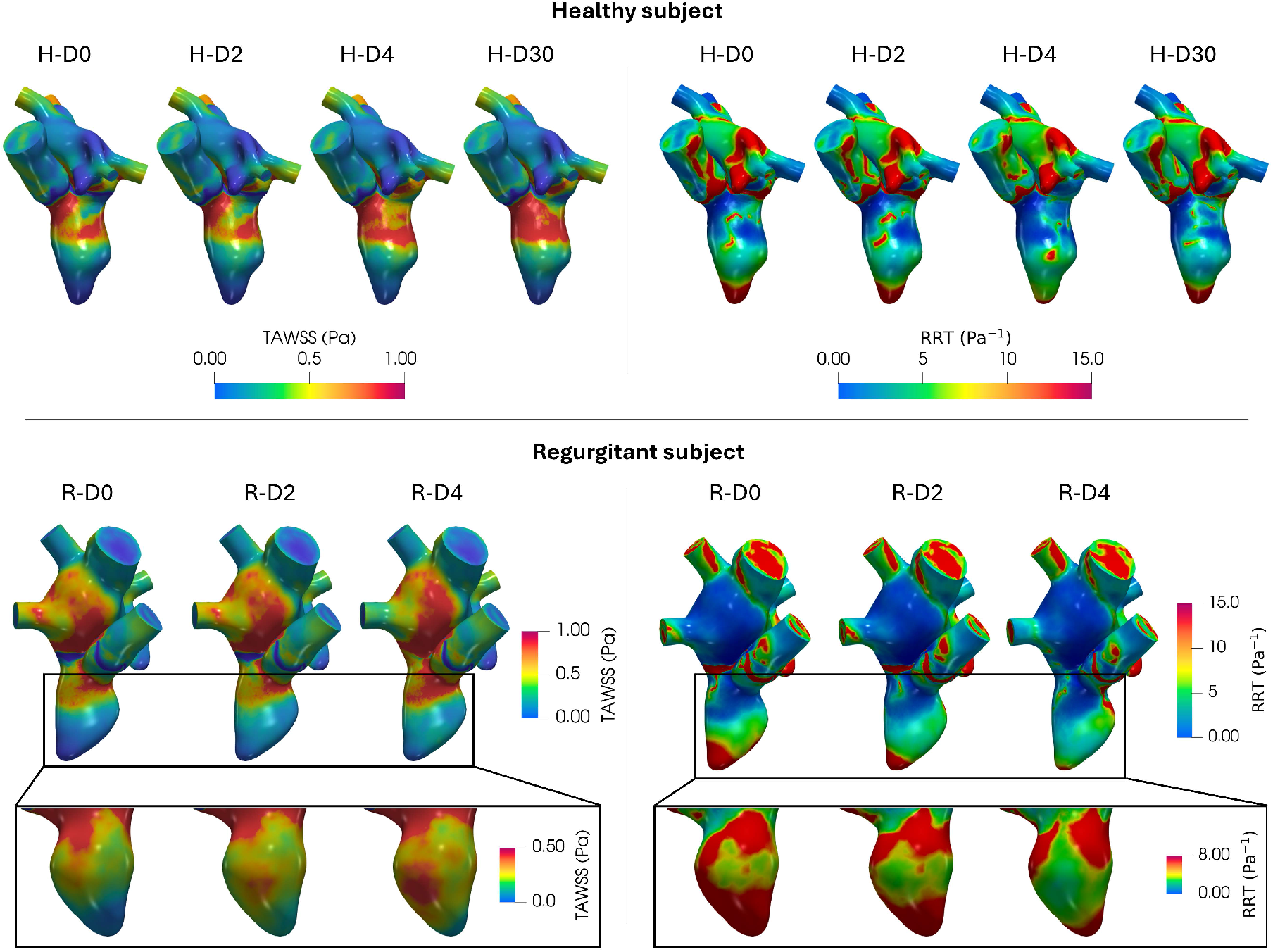
Top: Spatial distribution of TAWSS (left) and RRT (right) at the end-systolic configuration, for each scenario of healthy subject. Bottom: Spatial distribution of TAWSS (left) and RRT (right) at the end-systolic configuration, for each scenario of diseased patient.

#### 4.2.2 Regurgitant subject

This subsection describes the comparison results among the three different scenarios (R-D0, R-D2 and R-D4, refer to Table 1) in case of the diseased patient.

Figure 14 (left) shows the ensemble velocity magnitude computed in the regurgitant section (i.e., the plane where regurgitation assumes its maximum values in terms of velocity and spatial extension) at four time instants of the heartbeat. At systolic peak *t*_*PS*_ both the regurgitant jet and the velocity inside the ventricle feature similar patterns for all the scenarios. At the peak of E-wave *t*_*E*_, similarly to healthy case, two vortex rings develop below the two leaflets in all the scenarios. More in detail, *t*_*E*_ is the instant when the diastolic jet reaches the ventricular apex in R-D4 scenario; this happens later for R-D0 and R-D2. Moreover, in correspondence of the ventricular center and close to its apex, two vortexes develop for R-D2 and R-D4 scenarios, whereas in R-D0 they are absent. At the peak of diastasis *t*_*D*_ the mitral valve undergoes partial closure in R-D4, whereas in the remaining two cases it remains fully open. In R-D0 and R-D2 scenarios similar velocity patterns develop, with a coherent vortex structure in the middle of the ventricle, whereas in R-D4 scenario more chaotic eddies are present with localized larger velocities than the other two cases in correspondence of the anterior leaflet. At the peak of A-wave *t*_*A*_ R-D0 scenario still shows the vortex in the middle of the ventricle developed during the diastasis. Conversely, in R-D2 and R-D4 more swirling eddies are present. Moreover, R-D4 experiences higher velocities in correspondence of MV and below the anterior leaflet than the other two cases.

**Fig. 14.**
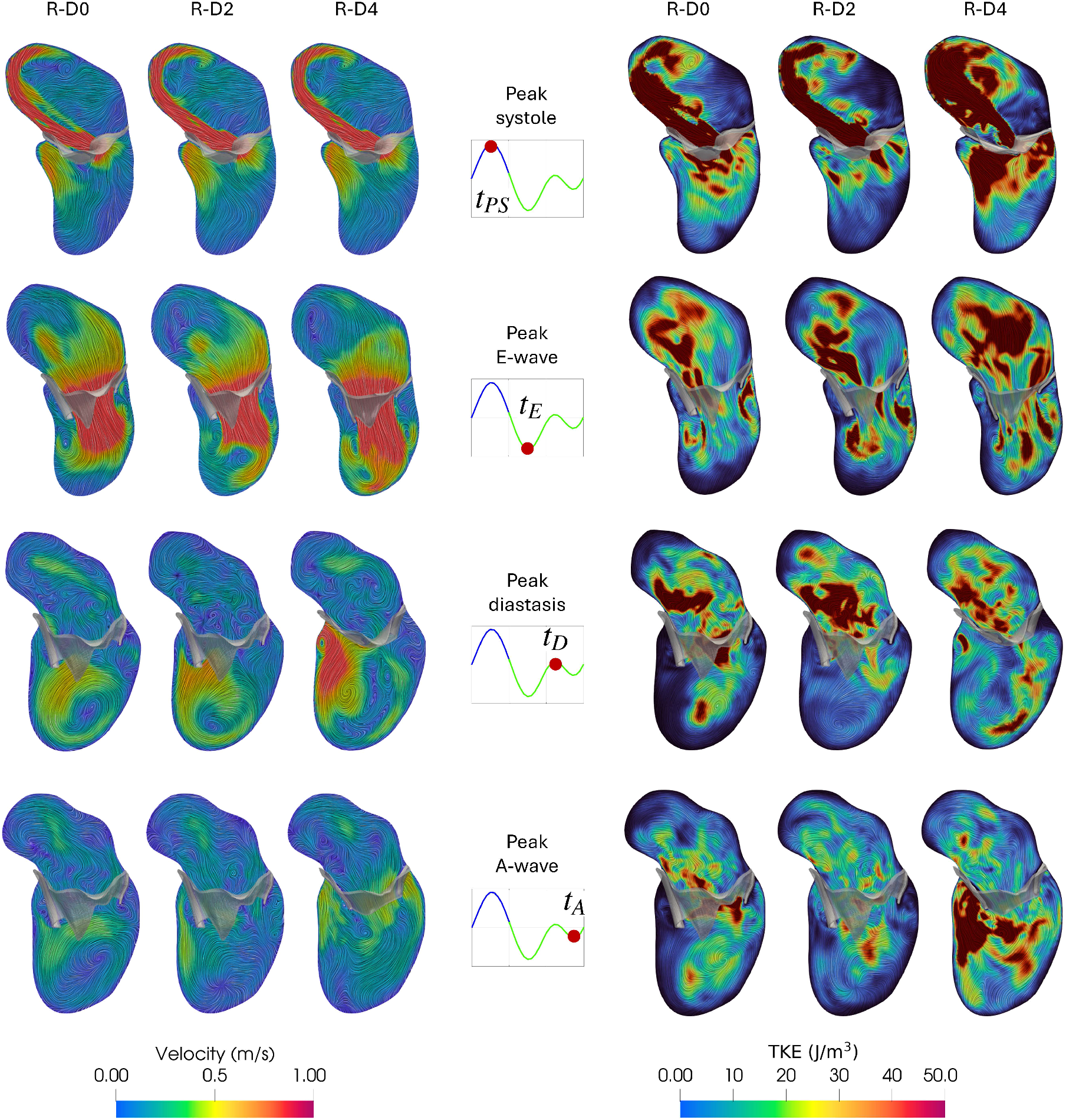
Magnitude of the ensemble velocity field (left) and spatial distribution of TKE (right) computed over five heartbeats at the peak of systole (*t*_*P S*_), peak of E-wave (*t*_*E*_), peak of diastasis (*t*_*D*_) and peak of A-wave (*t*_*A*_) in the three scenarios. The flow rate diagrams refer to the flow through the aortic (from 0 to 0.22 s, in blue) and the mitral orifice (from 0.22 to 0.8 s, in green). Regurgitant subject.

Figure 14 (right) shows the corresponding spatial distribution of TKE in the same section at the same instants. At *t*_*PS*_ the TKE distribution in atrium is similar in the three scenarios, with fluctuations pronounced especially in correspondence of the regurgitant jet. In particular, R-D4 scenario features a greater extension of high turbulence regions than R-D0 and R-D2, both in atrium and ventricle. At *t*_*E*_ all the scenarios experience a similar TKE pattern in the ventricle, with large values below the leaflets, in correspondence of the velocity vortexes observed at the corresponding instant (Figure 14, *t*_*E*_, left); furthermore, R-D4 presents also high values in the middle of the ventricle. Concerning the atrium, in R-D0 and R-D2 scenarios the TKE distribution is similar, with maximum values above the anterior leaflet, whereas R-D4 scenario features high values in a larger area, especially in the central region of the atrium. At *t*_*D*_ for all the scenarios the highest turbulence occurs in the atrium, especially in its middle region and close to the anterior leaflet. Concerning the ventricle, R-D4 scenario presents high TKE values along the ventricular wall close to the apex, while D2 exhibits almost no turbulence formation in the ventricle; instead, in R-D0 scenario, large values of TKE result in the middle of the ventricle. At *t*_*A*_ D4 features higher turbulence formation in the ventricle than R-D0 and R-D2 especially in the center of the ventricle and in LVOT region.

This TKE behavior has a consequence on GTKE time evolution. Considering the atrium (Figure 12, bottom-left), GTKE behavior is very similar between the three cases, with a peak in correspondence of *t*_*PS*_ (about 0.21 s), and coherent with other literature works [24] (where an average GTKE among different patients is obtained by means of phase-contrast MRI). Considering the ventricle (Figure 12, bottom-right), comparable values of GTKE are obtained until the instant of E-wave peak (about 0.45 s). Then, R-D4 features higher values with a maximum reached in correspondence of W-wave peak (about 0.7 s), see also Figure 14, right.

Figure 13 (bottom) shows the spatial distribution of the ensemble TAWSS and RRT, whose patterns are comparable for all the scenarios. Nevertheless, in R-D4 scenario elevated TAWSS values result along the ventricular wall, attributed to the different velocity patterns developed during diastole (see Figure 14, left). Consequently, R-D4 scenario exhibits lower RRT values in the ventricle compared to R-D0 and R-D2, suggesting that also in the patient with mitral valve regurgitation the inclusion of diastasis and A-wave could better describe the washout process developed by the left ventricle during diastole.

#### 4.2.3 Discussion on the choice of the mitral valve dynamic model in cardiac hemodynamics

This section aimed to analyze different models of mitral valve dynamics, in order to compare the fluid dynamic results in the context of DIB-CFD modeling. This is motivated by the fact that several studies in literature consider only some diastolic mitral valve configurations, while in other works [8, 7] all the configurations are considered.

Considering the different numerical results shown in previous Subsections for healthy and diseased cases (Subsection 4.2.1 and Subsection 4.2.2 respectively), it is possible to state that the inclusion of diastasis and A-wave configurations in mitral valve dynamics (D4 and D30 models) has important fluid dynamics consequences. Indeed, diastasis is characterized by the semi-closure of the mitral valve, which directs the flow towards the ventricular wall. This allows the blood to create a single, well-defined, clockwise vortex developing in the central region of the ventricle, facilitating the blood washout in the following systole. Moreover, modeling the diastasis allows to take into account the role that the mitral jet has in amplifying the shear stresses, especially in correspondence of the ventricular wall where the mitral valve directs the blood flow during diastasis. This results in a reduction of the residence time of blood particles, avoiding blood stagnation and reducing the heart work. Moreover, since the inclusion of diastasis implies a more pronounced rotational behavior of the blood, D4 and D30 MV models allows to better quantify the ventricular turbulence, both in terms of spatial distribution and of GTKE.

On the other hand, considering the fluid dynamics in atrium, no substantial difference between the scenarios is present, also in case of regurgitant heart. This is due to the fact that the different models of MV dynamics do not particularly influence the entity and the extension of regurgitant jet.

An important consideration concerns the comparison of D4 and D30 scenarios. We recall that this latter is the reference framework, since it is built reconstructing all the mitral valve geometries for all the available patient-specific images.

Our results reveal similar behaviors for almost the quantities analyzed, except for the turbulence at diastasis, where D30 features higher values. Furthermore, D4 model has the advantage of avoiding MV reconstruction at each frame, which are subjected to the operator sensitivity during the segmentation process and to a greater probability of noise, due to the acquisition protocol.

On the other hand, considering D0 and D2 models (thus, a reconstruction of only fully open and fully closed configurations) implies to neglect the MV dynamic during diastasis. Indeed, MV always remains totally open for the entire diastolic duration and multiple vortexes of medium intensity develop in the ventricles. Thus, these scenarios are less accurate in terms of ventricular turbulence and washout analysis. Moreover, since the results obtained in D0 and D2 scenarios are comparable, it is possible to conclude that providing a physiological time-imposed MV dynamic has no additional benefits compared to modeling instantaneous opening/closure of the mitral valve. Summarizing, we can state that in order to study in a more accurate way the diastolic dynamics of the mitral valve and its effects on the ventricular flow, an image-based MV dynamics should be considered, but reconstructing the valve only at significant temporal acquisitions.

## 5 Limitations and perspectives

The present study presents the following limitations:

1. Only one healthy and one diseased subjects are considered. This is a consequence of the sophisticated MV imaging acquisition protocol (not daily available), which is applied in order to build the different MV dynamic models;
2. Chordae tendineae are manually reconstructed due to the insufficient spatial resolution of the available imaging. Thus, the process is manual, so it is subjected to the operator sensitivity and this implies some limitations in reconstruction: a number of chordae equal for both subjects, the absence basal chordae, and the neglecting of bifurcations;
3. The absence of a unique global framework which is able to integrate both the subvalvular apparatus structures and a complex MV dynamics model in a single numerical experiment. This would require a huge computational effort, related to solving a very complicated RIIS problem, that accounts for both CT and MV dynamics. This will be considered in future studies;
4. In the case of sub-valvular simulations, we consider only one heartbeat and thus we are not able to be independent on the null initial conditions. As done for the study of Section 4, we need to extend to more heartbeat the analysis of the influence of chordae and papillaries;
5. For the aortic valve, we have a disposal only the open and closed configurations, thus only D0 and D2 models are considered. However, this is in our opinion acceptable, since the main focus in this paper is on the mitral valve dynamics and since no partial closure of the aortic valve (as happens for the mitral one during diastasis) is experienced during the systolic phase.

As a final remark, we stress that this paper try to shad a new light in the modeling of sub-valvular apparati and of mitral valve dynamics. Of course, future investigations will be needed to overcome the limitations of this work and to apply a comparison proposed in this work to more patients and diseased cases of clinical relevance.

## Acknowledgements

AC and CV are members of the INdAM group GNCS “Gruppo Nazionale per il Calcolo Scientifico” (National Group for Scientific Computing). CV has been partially supported by the Italian Ministry of University and Research (MIUR) within the PRIN (Research projects of relevant national interest) MIUR PRIN22-PNRR n. P20223KSS2 “Machine learning for fluid-structure interaction in cardiovascular problems: efficient solutions, model reduction, inverse problems”, and by the Italian Ministry of Health within the PNC PROGETTO HUB LIFE SCIENCE - DIAGNOSTICA AVANZATA (HLS-DA) “INNOVA”, PNC-E3-2022-23683266?CUP: D43C22004930001, within the “Piano Nazionale Complementare Ecosistema Innovativo della Salute” - Codice univoco investimento: PNC-E3-2022-23683266.

## Authors contributions

Conceptualization: L. Bennati, C.Vergara.

Methodology: A. Crispino, L. Bennati, C.Vergara. Numerical experiments: A. Crispino, L. Bennati. Formal analysis and investigation: A. Crispino, L. Bennati.

Writing - original draft preparation: A. Crispino, C.Vergara.

Writing - review and editing: L. Bennati. Funding acquisition: C.Vergara.

Resources: A. Crispino, L. Bennati, C.Vergara. Supervision: C.Vergara.

## Ethical statements

Ethical Review Board of Borgo Trento Hospital, Verona approval and informed consent were obtained from all patients.

